# Uncovering hidden assembly artifacts: when unitigs are not safe and bidirected graphs are not helpful

**DOI:** 10.1101/2022.01.20.477068

**Authors:** Amatur Rahman, Paul Medvedev

## Abstract

Recent assemblies by the T2T and VGP consortia have achieved significant accuracy but required a tremendous amount of effort and resources. More typical assembly efforts, on the other hand, still suffer both from mis-assemblies (joining sequences that should not be adjacent) and from under-assemblies (not joining sequences that should be adjacent). To better understand the common algorithm-driven causes of these limitations, we investigated the unitig algorithm, which is a core algorithm at the heart of most assemblers. We prove that, contrary to popular belief, even when there are no sequencing errors, unitigs are not always safe (i.e. they are not guaranteed to be substrings of the sequenced genome). We also prove that the unitigs of a bidirected de Bruijn graph are different from those of a doubled de Bruijn graph and, contrary to our expectations, result in under-assembly. Using experimental simulations, we then confirm that these two artifacts exist not only in theory but also in the output of widely used assemblers. In particular, when coverage is low then even error-free data results in unsafe unitigs; also, unitigs may unnecessarily split palindromes in half if special care is not taken. To the best of our knowledge, this paper is the first to theoretically predict the existence of these assembler artifacts and confirm and measure the extent of their occurrence in practice.

## 1 Introduction

Reconstructing the full sequence of a genome from its sequencing data remains one of the most challenging problems in bioinformatics. Assemblers have suffered both from mis-assemblies (putting together sequences that should not be adjacent) and under-assemblies (not putting together sequences whose adjacency should be apparent from the data) [4, 35]. Recent efforts by the Telomere-to-Telomere consortium [25, 24] and the Vertebrate Genome Project [30] demonstrated how long read technologies, long-range contact mapping, and manual curation can alleviate these errors. However, the time and cost of those efforts remain prohibitive for most biology labs. In such cases, mis- and under-assemblies continue to be a major limitation (e.g. [38]).

Understanding the common algorithm-driven causes of these limitations is made complicated by the diversity and complexity of assembly algorithms. We can start by focusing on assemblers that use de Bruijn graphs (dBGs) [16], which continue to be popular even for long-read data [5]. But even dBG-based assemblers differ on how they handle complexities arising from sequencing errors, heterogeneity, or DNA double strandedness. Nevertheless, most assemblers are built on top of the unitig algorithm, which returns all the maximal unitigs in an assembly graph [35]; a unitig is a path whose vertices have exactly one incoming and one outgoing edge, with the exception that the first and last vertex can have any number of incoming and outgoing edges, respectively. Being a common denominator of most assemblers, the unitig algorithm is a good target for investigating shared sources of mis- and under-assemblies.

It is already known that the unitig algorithm contributes to under-assembly (e.g. see the safe and complete framework of [37, 9]) and can trivially create mis-assemblies when there are sequencing errors. The effect of sequencing errors on assembly errors has even been theoretically studied more broadly in [32, 33, 34]. However, it is widely assumed that if it were not for sequencing errors, unitigs would always be safe (i.e. substrings of the sequenced genome). In an earlier work [19], we attempted to formally prove this but could only do so by assuming perfect coverage. This assumption was also necessary in another earlier work [37], where it was suggested that without it, unitigs may not be safe. Unitigs were also implied to be unsafe in certain models of the assembly problem [9]. We therefore hypothesize that, contrary to popular belief, there are non-contrived conditions which lead to unsafe unitigs on error-free data.

The unitig algorithm also needs to account for the fact that the strand from which a read is sequenced is unknown. Most assemblers do so via two common approaches to constructing the dBG. In one, every *k*-mer is “doubled” prior to constructing the dBG, i.e. for every *k*-mer in the input, both it and its reverse complement is added to the dBG (e.g. SPAdes [6]). In the other approach, edges are given two instead of one orientation, thereby capturing the way that double-stranded strings can overlap. This results in a bidirected dBG [22], used in assemblers such as ABySS [36, 17]). Since this is a more elegant construction for capturing the double-stranded nature of the data, one would intuitively expect that it should not hurt assembly accuracy.

In this paper, we perform a theoretical and empirical study to validate our two hypothesis about common algorithm-driven sources of mis- and under-assemblies. First, despite widespread belief to the contrary, we show that even on error-free data, unitigs do not always appear in the sequenced genome (i.e. they are *unsafe*). Our experimental results confirm that at least two different assemblers exhibit this behavior in practice. Second, we establish that there is a bijection between maximal unitigs in the doubled and bidirected dBGs, except that palindromic unitigs in the doubled dBG are split in half in the bidirected dBG. This shows that, contrary to intuition, naively using the bidirected graph actually contributes to under-assembly compared to the doubled graph. Our experimental results confirm that this artifact appears in some assemblers but not in others. Nevertheless, we also find that the extent of these two artifacts is limited. To the best of our knowledge, this paper is the first to theoretically predict the existence of these assembler artifacts and confirm and measure the extent of their occurrence in practice.

## 2 Preliminaries

In this section we give the formal definitions for our paper. The reader may wish to delay reading the last three paragraphs (relating to bidirected graphs) until they are used in Section 4.

### Strings

In this paper, we assume all strings are over the four-letter DNA alphabet. A string of length *k* is called a *k-mer*. We define *suf*_*k*_(*x*) (respectively, *pre*_*k*_(*x*)) to be the last (respectively, first) *k* characters of *x*. When the subscript is omitted from *pre*, and *suf*, we assume it is *k* − 1. For *x* and *y* with *suf* (*x*) = *pre*(*y*), we define *gluing x* and *y*, denoted by *x* ⊙ *y*, as *x* concatenated with the last |*y*| − *k* + 1 characters of *y*. Given two strings *x* and *y*, we define *occ*_*y*_(*x*) as the number of times that *x* occurs in *y*. The reverse complement of *x* is denoted as 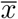. For a set of strings *𝒮*, 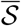 denotes the set of the reverse complements of all strings of *𝒮*. A string *x* is a *palindrome* iff 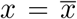. A string *x* is *canonical* if it is the lexicographically smaller of *x* and 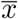. For *s* ∈ {0, 1}, we define *orient*(*x, s*) to be *x* if *s* = 0 and to be 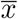 if *s* = 1. To *canonize x* is to replace it by its canonical version, *canon*(*x*) = min_*i*_(*orient*(*x, i*)). We say that *x*_0_ and *x*_1_ have a (*s*_0_, *s*_1_)*-oriented-overlap* if *suf* (*orient*(*x*_0_, 1 − *s*_0_)) = *pre*(*orient*(*x*_1_, *s*_1_)). Intuitively, such an overlap exists between two strings if we can orient them in such a way that they are glueable. For example, *GTT* and *TTG* has a (1, 0)-oriented overlap, and *AAC* and *TTG* have a (0, 0)-oriented overlap. We define the *non-canonical k-spectrum* sp^*k*^(*x*) as the set of all *k*-mer substrings of *x*.

### Directed de Bruijn graphs

Given a set of *k*-mers *K*, the *basic node-centric directed de Bruijn Graph, G*_basic_(*K*), is directed graph where nodes are the *k*-mers of *K*, and an edge exists from *k*-mer *x* to *k*-mer *y* iff *suf* (*x*) = *pre*(*y*). A *double directed de Bruijn graph* on *K, G*_dbl_(*K*) is a basic de Bruijn graph on the set of *k*-mers 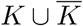, i.e. 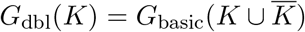. Observe that for any *k*-mer *x* such that 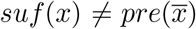, the existence of the edge from *x* to *y* in *G*_dbl_(*K*) implies the existence of a different edge from 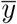 to 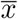. We refer to such a pair of edges as *mirrors*. For a *k*-mer *x* such that 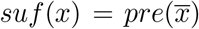, the *G*_dbl_(*K*) will contain an edge from *x* to *x*; we call this edge a *self-mirror*.

### Walks and unitigs in directed graphs

For a vertex *x* in a directed graph, the *in-degree d*^−^(*x*) (respectively, *out-degree d*^+^(*x*)) is the number of edges incoming to (respectively, outgoing from) it. The sequence of vertices *w* = (*x*_0_, …, *x*_*n*_), for *n* ≥ 0, is a *walk* iff for all 1 ≤ *i* ≤ *n* there exists an edge from *x*_*i* −1_ to *x*_*i*_. Vertices *x*_0_ and *x*_*n*_ are called *endpoints*, and a walk sometimes has one endpoint. The *spelling* of a walk is defined as *spell*(*w*) = *x*_0_ ⊙· · · ⊙ *x*_*n*_. A walk is said to be *circular* iff *n* ≥ 1 and *x*_0_ = *x*_*n*_, and a simple *cycle* if for all *i* and *j* such that 0 ≤ *i < j* ≤ *n, x*_*i*_ = *x*_*j*_ implies *i* = 0 and *j* = *n*. A simple periodic cycle is a walk that starts with a simple cycle and then keeps on looping around it without ever exiting; formally, a walk is a *simple periodic cycle* if there exists 0 ≤ *i* ≤ *n* − 1 such that (*x*_0_, …, *x*_*i*_) is a simple cycle and *x*_*i*+1_, …, *x*_*n*_ is a repetition of *x*_0_, …, *x*_*i*_, except the last repetition may be partial. A walk is a *unitig* if it is not a periodic cycle and for all 1 ≤ *i* ≤ *n, d*^−^(*x*_*i*_) = 1 and for all 0 ≤ *i* ≤ *n* − 1, *d*^+^(*x*_*i*_) = 1. A unitig is *maximal* if it is not a proper subwalk of another unitig.

### Bidirected de Bruijn graph

A *bidirected graph G* is a pair (*V, E*) where the set *V* are called vertices and *E* is a set of edges. Intuitively, every vertex has two sides; formally, a *vertex-side* is a pair (*u, s*), where *u* ∈ *V* and *s* ∈ {0, 1}. An edge *e* is a set of two vertex-sides {(*u*_0_, *s*_0_), (*u*_1_, *s*_1_)}, where *u*_*i*_ ∈ *V* and *s*_*i*_ ∈ {0, 1}, for *i* ∈ {0, 1}. Intuitively, an edge is an undirected connection between two (not-necessarily distinct) vertex-sides. We say that an edge *e* is incident to each of the two vertex-sides. Note that there can be multiple edges between two vertices, but only one edge once the sides are fixed. A *labeled bidirected graph* is a bidirected graph *G* where every vertex *u* has a string label *lab*(*u*), and for every edge *e* = {(*u*_0_, *s*_0_), (*u*_1_, *s*_1_)}, there is a (*s*_0_, *s*_1_)-oriented-overlap between *lab*(*u*_0_) and *lab*(*u*_1_). *G* is said to be *overlap-closed* if there is an edge for every such overlap. Let *K* be a set of canonical *k*-mers. The node-centric *bidirected de Bruijn graph*, denoted by *G*_bid_(*K*), is the overlap-closed labeled bidirected graph where the vertices and their labels correspond to *K*. Figure S1A shows an example of a bidirected graph.

### Walks and unitigs in bidirected graphs

An edge in a bidirected graph is an *inverted loop* if its two vertex-sides are equal. An inverted loop {(*u, s*), (*u, s*)} is *lonely* if it is the only edge incident to (*u, s*). We define the *degree* of a vertex-side *d*(*u, s*) to be the number of edges incident to it, but with an inverted loop contributing two to the degree. A sequence *t* = (*u*_0_, *s*_0_, *u*_1_, *s*_1_, …, *u*_*n*_, *s*_*n*_) with *n* ≥ 0 is a *walk* if for all 1 ≤ *i* ≤ *n*, there exists an edge *e*_*i*_ = {(*u*_*i* −1_, 1 − *s*_*i* −1_), (*u*_*i*_, *s*_*i*_)}. The vertex-sides (*u*_0_, *s*_0_) and (*u*_*n*_, 1 − *s*_*n*_) are called the first and last *endpoint sides*, respectively. Note that even when *n* = 0, there are two endpoint sides. The *spelling* of a walk is defined as *spell*(*w*) = *orient*(*lab*(*u*_0_), *s*_0_) ⊙· · · ⊙ *orient*(*lab*(*u*_*n*_), *s*_*n*_). The reverse of *t* is *rev*(*t*) = (*u*_*n*_, 1 − *s*_*n*_, …, *u*_0_, 1 − *s*_0_). Note that, as expected, 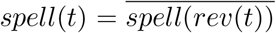. Note that if *t*′ is a subwalk of *t*, then *rev*(*t*′) is a subwalk of *rev*(*t*) and *spell*(*t*′) is a substring of *spell*(*t*) (the converse is not necessarily true when *k* is even). Figure S1BC gives an example illustrating a walk in a bidirected graph and Figure S1D shows a corresponding walk in a doubled directed dBG.

A walk *w* = (*u*_0_, *s*_0_, …, *u*_*n*_, *s*_*n*_) is said to be *circular* iff *n* ≥ 1 and (*u*_0_, *s*_0_) = (*u*_*n*_, *s*_*n*_), and a simple *cycle* if for all *i* and *j* such that 0 ≤ *i < j* ≤ *n*, (*u*_*i*_, *s*_*i*_) = (*u*_*j*_, *s*_*j*_) implies *i* = 0 and *j* = *n*. A *simple periodic cycle* is a walk that starts with a simple cycle and then keeps on looping around it without ever exiting; formally, *w* is a *simple periodic cycle* if there exists 0 ≤ *i* ≤ *n* − 1 such that (*u*_0_, *s*_0_, …, *u*_*i*_, *s*_*i*_) is a simple cycle and (*u*_*i*+1_, *s*_*i*+1_, …, *u*_*n*_, *s*_*n*_) is a repetition of (*u*_0_, *s*_0_, …, *u*_*i*_, *s*_*i*_), except the last repetition may be partial. A walk is a *unitig* if it is not a periodic cycle and for all 1 ≤ *i* ≤ *n, d*^−^(*x*_*i*_) = 1 and for all 0 ≤ *i* ≤ *n* − 1, *d*^+^(*x*_*i*_) = 1. A walk (*u*_0_, *s*_0_, …, *u*_*n*_, *s*_*n*_) is a *unitig* if it is not a periodic cycle and for all 0 ≤ *i < n, d*(*u*_*i*_, 1 − *s*_*i*_) = 1 and, for all 0 *< i* ≤ *n, d*(*u*_*i*_, *s*_*i*_) = 1. A unitig is said to be *maximal* if it is not a proper subwalk of another unitig. Note that all the subwalks of a unitig must themselves be unitigs.

## 3 Safety of unitigs

In this section, we will give necessary and sufficient conditions for a unitig to be unsafe in the basic dBG constructed from error-free reads. To properly formulate this question, we define a *sequenced read interval* as a genomic interval that generated a read, i.e. from which a read was sequenced. A sequencing experiment then corresponds to a set of sequenced read intervals. A *sequenced interval* is then defined as a maximal interval which is covered by sequenced read intervals, with the additional constraint that any two consecutive sequenced intervals overlap by at least *k* − 1. We define a *sequenced segment* as the string corresponding to a sequenced interval. Observe that the sequenced intervals do not overlap by more than *k* − 2 bases (otherwise they would not be maximal), but the sequenced segment may have longer overlaps due to repeats. A set of reads then induces a set *𝒮* = {*S*_1_, …, *S*_|*𝒮*|_} of sequenced segments. Figure 1 illustrates this formulation. In this section, we do not explicitly account for reverse complements, since they will be considered in Section 4.

**Figure 1:**
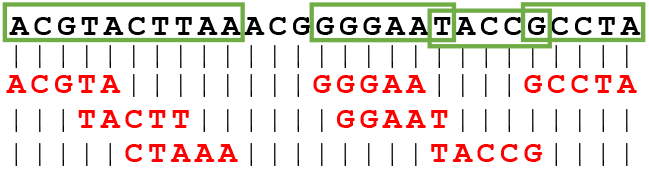
Illustration of sequenced segments. The black text on top shows the reference genome of length 26. The seven sequences in red are reads aligned to the reference. The green boxes highlight the resulting sequenced segments when *k* = 3. Note that the reads TACCG and GCCTA form two separate segments as the *k*-mer CGC is not present in *K*.

Given a set of sequenced segments *𝒮*, we say that a unitig *w* in *G*_basic_(sp^*k*^(*𝒮*)) is *unsafe* iff *spell*(*w*) is not a substring of a string in *𝒮*. Equivalently, *w* is unsafe iff it is not a subwalk of a walk that corresponds to a string in *𝒮*. Our definition of unsafe captures the notion of a potential mis-assembly, as the unitig is not present in the sequenced part of the genome.^1^ Observe that in formulating the problem, we start with the set of sequenced segments themselves; the read set that induced them is irrelevant. We can now state the main result of this section, which gives the necessary and sufficient conditions for a unitig to be unsafe. The proof of this theorem, along with the necessary Lemmas, is left for Appendix A due to space constraints.

### Theorem 1.

*Let 𝒮 be a set of sequenced segments and let w* = (*x*_0_, …, *x*_*m*_) *be a unitig in G*_*basic*_(*sp*^*k*^(*𝒮*)). *Then w is unsafe if and only if for all S* ∈ *𝒮, one of the following holds:*

i. *S does not contain any k-mer of w*,
ii. *occ*_*S*_(*pre*_*k*_(*S*)) = 1 *and pre*_*k*_(*S*) = *x*_*i*_ *for some* 1 ≤ *i* ≤ *m*,
iii. *occ*_*S*_(*suf*_*k*_(*S*)) = 1 *and suf*_*k*_(*S*) = *x*_*j*_ *for some* 0 ≤ *j* ≤ *m* − 1, *or*
iv. *occ*_*S*_(*pre*_*k*_(*S*)) = *occ*_*S*_(*suf*_*k*_(*S*)) = 2 *and there exists* 1 ≤ *i* ≤ *j* ≤ *m* − 1 *such that pre*_*k*_(*S*) = *x*_*i*_ *and suf*_*k*_(*S*) = *x*_*j*_.

The cases of Theorem 1 are illustrated in Figure 2 and can be understood intuitively as follows. Since every *k*-mer of *G*_basic_(sp^*k*^(*𝒮*)) is in *𝒮*, every *k*-mer of *w* must be touched by some *S* ∈ *𝒮*. Then, consider a walk *g* corresponding to such a string *S*. If *g* starts in the middle of *w* and does not visit its own starting vertex again, then *g* does not fully contain *w* (case (*ii*)). Similarly, if *g* ends in the middle of *w* and did not visit its own ending vertex previously, then *g* does not fully contain *w* (case (*iii*)). If *g* starts and ends in the middle of *w*, with the ending vertex to the right of the starting vertex, and contains each of those vertices exactly twice, then *g* does not fully contain *w* (case (*iv*)). This is the “if” direction of Theorem 1, with the “only if” direction further stating that under all other conditions, *g* fully contains *w*.

**Figure 2:**
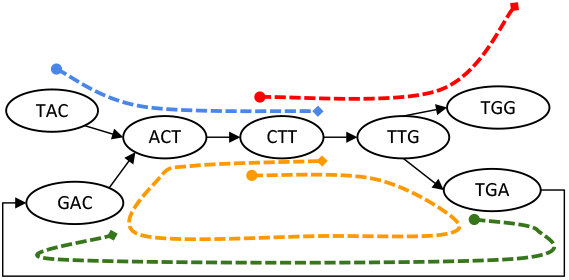
Illustration of the cases in Theorem 1. The graph in the figure represents *G*_basic_(sp^*k*^(*𝒮*)), where *k* = 3 and *𝒮* = {CTTGG, CTTGACTT, TACTT, TGAC}. The segments are marked by dashed lines with their starts marked with a dot and their ends marked with a diamond. The unitig *w* = {*ACT, CT T, TT G*} is un-safe because for each of the segment, one of the cases in Theorem 1 is true. For segment colored in green, case (i) holds. For red, case (ii), for blue, case (iii) and for orange, case (iv) holds.

When the genome is a single chromosome and the coverage is high enough so that every *k*-mer is sequenced, the whole genome becomes one sequenced segment. In this case, Theorem 1 simplifies because the genome has only one starting and ending vertex and, for a unitig *w* to be unsafe, the genome must somehow contain every vertex of *w* without containing *w* as a subwalk.

### Corollary 1.

*Let X be a string and let w* = (*x*_0_, …, *x*_*m*_) *be a unitig in G*_*basic*_(*sp*^*k*^(*X*)). *Then spell*(*w*) *is not a substring of X iff one of the following holds:*

1. *occ*_*X*_(*pre*_*k*_(*X*)) = *occ*_*X*_(*suf*_*k*_(*X*)) = 1, *pre*_*k*_(*X*) = *x*_*i*_, *suf*_*k*_(*X*) = *x*_*i* −1_ *for some* 1 ≤ *i* ≤ *m*.
2. *occ*_*X*_(*pre*_*k*_(*X*)) = *occ*_*X*_(*suf*_*k*_(*X*)) = 2, *pre*_*k*_(*X*) = *x*_*i*_, *suf*_*k*_(*X*) = *x*_*j*_ *for some* 0 *< i* ≤ *j < m.* *Moreover, this can hold for at most one unitig in G*_*basic*_(*sp*^*k*^(*X*)).

This corollary tells us that with perfect coverage, all unitigs, except possibly one, are safe. Note that this is a stronger version of the perfect coverage case than the one given in [19], which made an assumption that the starting vertex of *X* is a source and the ending vertex of *X* is a sink.

A natural question is how a scenario which gives an unsafe unitig looks like in terms of the original genome. Figure 3 visualized the following natural possibility. Suppose that the sequenced genome *X* has a repeat that appears as a maximal unitig ψ in *G*_basic_(sp^*k*^(*X*)). Then, suppose that the region encompassing the start of one copy and the region encompassing the end of the other copy is not sequenced. Then ψ loses its maximality in *G*_basic_(sp^*k*^(*𝒮*)) and becomes a subwalk of a bigger unitig *w*. Though *w* is a unitig in the graph from the sequencing data, it would not be a unitig if all the *k*-mers of *X* were included in the graph. In Section 5, we will show that this situation accounts for the majority of our experimental observations.

**Figure 3:**
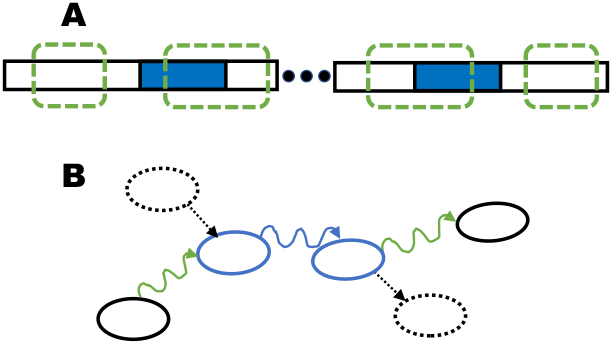
Panel A shows two parts of a sequenced genome *X*. Regions surrounded by green dashed boxes are the sequenced segments *𝒮*. The solid blue boxes represent two copies of a repeat. Panel B shows the resulting *G*_basic_(sp^*k*^(*𝒮*)), with dashed vertices and edges representing vertices that are in *G*_basic_(sp^*k*^(*X*)) but not sequenced.

## 4 The relationship between the doubled dBG (*G*_dbl_(*K*)) and the bidirected dBG (*G*_bid_(*K*))

In this section, we will characterize the relationship between the maximal unitigs of *G*_dbl_(*K*) and the maximal unitigs of *G*_bid_(*K*) (Theorem 2). Due to space constraints, the lemmas and proofs needed to prove Theorem 2 are in Appendix B. Here, we will instead give an intuitive walk-through to elucidate the relationship between the two graphs. We will incrementally show the relationship between objects in the doubled graph and the bidirected graph — first between vertices and vertex-sides, then between edges, then between walks, and finally between maximal unitigs.

Let *K* be a set of canonical *k*-mers, with *k* odd. We only consider the case of odd *k*; when *k* is even, there may be palindrome *k*-mers, which create special cases to handle both in the practical assembler implementation and in the theoretical analysis. Since most assemblers anyway restrict *k* to be odd, we limit ourselves to this case as well.

There is a natural mapping between vertices of *G*_dbl_(*K*) and vertex-sides of *G*_bid_(*K*). For a vertex *x* in *G*_dbl_(*K*), define *F*_*V*_ (*x*) = (*u, s*), where *u* is a vertex in *G*_bid_(*K*) and *s* ∈ {0, 1} such that *lab*(*u*) = *orient*(*x, s*). By the definition of *G*_bid_(*K*), there exists a unique *u* and unique *s* that satisfy this condition. The uniqueness of *s* is guaranteed by the fact that *x* cannot be a palindrome. Formally, *F*_*V*_ is a bijection between vertices of *G*_dbl_(*K*) and vertex-sides of *G*_bid_(*K*) (Lemma B.10). There is also a natural mapping between edges in *G*_dbl_(*K*) and *G*_bid_(*K*). Let *x*_1_ and *x*_2_ be two *k*-mers in *G*_dbl_(*K*) and let (*u*_1_, *s*_1_) = *F*_*V*_ (*x*_1_) and (*u*_2_, *s*_2_) = *F*_*V*_ (*x*_2_). We define the mapping

*F*_*E*_(*x*_1_, *x*_2_) = {(*u*_1_, 1 − *s*_1_), (*u*_2_, *s*_2_)} such that (*x*_1_, *x*_2_) is an edge in *G*_dbl_(*K*) if and only if *F*_*E*_(*x*_1_, *x*_2_) is an edge in *G*_bid_(*K*) (Lemma B.11). Note, however, that *F*_*E*_ is not a bijection, since a pair of mirror edges (*x, y*) and 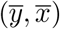 map to the same bidirected edge, i.e. 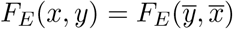.

The *F*_*V*_ and *F*_*E*_ mappings allow us to naturally define a mapping from walks in *G*_dbl_(*K*) to walks in *G*_bid_(*K*). Let *w* = (*x*_0_, …, *x*_*n*_) be a walk in *G*_dbl_(*K*). For each 0 ≤ *i* ≤ *n*, let (*u*_*i*_, *s*_*i*_) = *F*_*V*_ (*x*_*i*_) and define *F*_*W*_ (*w*) ≜ (*u*_0_, *s*_0_, …, *u*_*n*_, *s*_*n*_). *F*_*W*_ is a spell-preserving bijection between the set of walks in *G*_dbl_(*K*) and the set of walks in *G*_bid_(*K*) (Lemma B.12).

One might hypothesize that *F*_*W*_ is also a bijection between the maximal unitigs of *G*_dbl_(*K*) and the maximal unitigs of *G*_bid_(*K*). Surprisingly, it turns out to not be the case, though the following more careful analysis reveals a close relationship. For *G*_dbl_(*K*), let us partition the set of maximal unitigs into non-palindromic strings *D*_non-pal_ and palindromic strings *D*_pal_. For *G*_bid_(*K*), let *B*_no-loop_ be the set of maximal unitigs where neither endpoint side has an incident lonely inverted loop, let *B*_first-loop_ be the set of maximal unitigs where the only endpoint side with a lonely inverted loop is the first one, and let *B*_last-loop_ be the set of maximal unitigs where the only endpoint side with a lonely inverted loop is the last one. To avoid corner cases, let us further assume that there are no circular unitigs in *G*_dbl_(*K*), which eliminates the possibility of a maximal unitig having lonely inverted loops at both endpoint sides and implies that *B*_no-loop_, *B*_first-loop_, and *B*_last-loop_ are a partition of the maximal unitigs of *G*_bid_(*K*) (Lemma B.16). Figure 4 shows an example.

**Figure 4:**
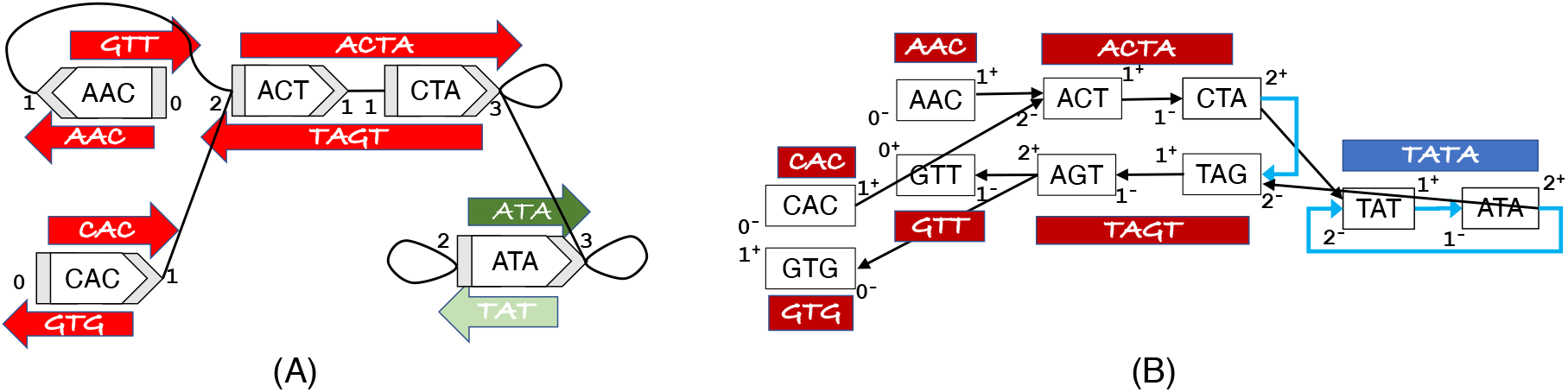
Example of a bidirected dBG (*G*_bid_(*K*)) (panel A) and a doubled dBG (*G*_dbl_(*K*)) (panel B) on the same underlying set of *k*-mers *K* = {*CAC, AAC, ACT, CT A, ATA*}. Each vertex side in *G*_bid_(*K*) and each in- and outside of a vertex in *G*_dbl_(*K*) is numbered with the corresponding degree. All maximal unitigs are shown using a long filled rectangle with an arrow. The maximal unitigs of *G*_bid_(*K*) are color coded so that red is *B*_no-loop_, dark green is *B*_last-loop_, and light green is *B*_first-loop_. The maximal unitigs of *G*_dbl_(*K*) are color coded so that dark red is *D*_non-pal_ and blue is *D*_pal_. Self-mirror edges in *G*_dbl_(*K*) are shown in blue.

We also need to define a function head which, informally, takes a maximal palindromic unitig in *G*_dbl_(*K*), extracts the first half of it, and maps it to *G*_bid_(*K*). Formally, head(*w*) maps a walk *w* = (*x*_0_, …, *x*_*n*_) in *D*_pal_ to the walk 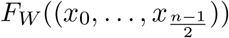 in *G*_bid_(*K*). Note that 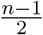 is necessarily an integer since *w* is a palindrome and hence *n* must be odd (Lemma B.1). We can now state the main theorem of this section.

### Theorem 2.

*Let K be a set of canonical k-mers where k is odd and G*_*dbl*_(*K*) *does not contain a circular unitig*.

i. *The function F*_*W*_ *is a bijection from D*_*non-pal*_ *to B*_*no-loop*_.
ii. *The function rev is a bijection between B*_*last-loop*_ *and B*_*first-loop*_.
iii. head *is a bijection from D*_*pal*_ *and B*_*last-loop*_

Figure 5 schematically illustrates the relationship captured by Theorem 2. The theorem says that for maximal unitigs that are non-palindromic in *G*_dbl_(*K*) and do not have inverted self loops incident at the endpoint sides in *G*_bid_(*K*), *F*_*W*_ is in fact a bijection. However, every maximal unitig *w* that is palindromic in *G*_dbl_(*K*) is split into two maximal unitigs in *G*_bid_(*K*): one that spells the first half of *w* and has a self loop incident at the last endpoint side, and one that spells the second half of *w* and has a self loop at the first endpoint side. These are necessarily reverses of each other.

**Figure 5:**
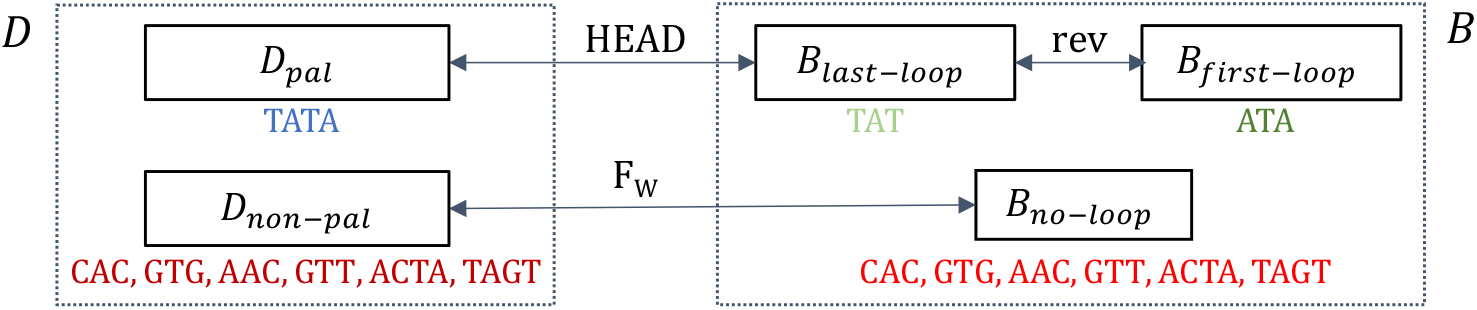
Overview of relationship between maximal unitigs in double and bidirected graph for odd *k*. We use the example from Figure 4, where *K* = {*AAC, ACT, CT A, CAC, ATA*}. The set of maximal unitigs from *G*_dbl_(*K*), *D* is partitioned into *D*_pal_ and *D*_non-pal_. The set of maximal unitigs from *G*_bid_(*K*), *B* is partitioned into *B*_last-loop_, *B*_first-loop_ and *B*_no-loop_. The arrows between different subsets of *D* and *B* denote bijections.

Inverted loops are caused by *k*-mers *x* where 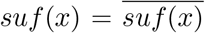 (e.g. GTA). When these type of *k*-mers are not present in *K*, there are no inverted loops in *G*_bid_(*K*) or palindromic unitigs in *G*_dbl_(*K*). Hence, *D*_pal_ = *B*_first-loop_ = *B*_last-loop_ = ∅, and Theorem 2 immediately simplifies.

### Corollary 2.

*Let K be a set of k-mers, with odd k, which does not contain any x such that* 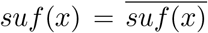. *Then F*_*W*_ *is a bijection from the maximal unitigs in G*_*dbl*_(*K*) *to the maximal unitigs in G*_*bid*_(*K*).

## 5 Empirical results

### Occurrence of unsafe unitigs in real genomes

Theorem 1 predicts the possibility of unsafe unitigs. To verify the extent to which this happens with real genomes, we use T2T human reference chromosome 1 [25]. We simulated error-free reads of length 100 with varying target coverages and varying *k*. Note that for this experiment, we want to test if mis-assemblies occur even when the data is perfect, so making the reads error-free is necessary. The sequenced read intervals correspond to the source location of each simulated read, and the sequenced segments are defined as in Section 3. From these reads, we constructed the basic de Bruijn graph and output its maximal unitigs, using a version of BCALM [10, 11] modified to ignore reverse-complementary. We confirmed that the unitigs that were unsafe(i.e. not a substring of the sequenced segments) were exactly the unitigs that satisfied the conditions of Theorem 1.

Table 1 shows the number of unsafe unitigs, as a function of the coverage and of *k*. There are as many as 17,635 unsafe unitigs (at coverage 2x and *k* = 71). The best indicator for the number of unsafe unitigs is the percent of *k*-mers sampled (or the number of sequenced segments), i.e. the number of unsafe unitigs goes down as the percent of sampled *k*-mers goes up. This trend is in line with the prediction of Corollary 1, which states that once the coverage is perfect, we expect to see at most one unsafe unitig. Our results indicate that the artifacts identified by Theorem 1 do occur in real genomes, though they become less common as more of the genomic *k*-mers are sampled.

**Table 1:** The presence of unsafe and mis-assembled unitigs in human chromosome 1, using simulated error-free reads.

| coverage | $k$ | % $k$-mers sampled | n. sequenced segments | n. unitigs | n. unsafe | n. mis-assembled | n. Figure 3 cases |
| --- | --- | --- | --- | --- | --- | --- | --- |
| 1x | 71 | 26.47 | 1,838,685 | 1,747,456 | 12,396 | 449 | 383 |
| 2x |  | 45.94 | 2,710,240 | 2,582,737 | 17,635 | 708 | 628 |
| 10x |  | 95.82 | 1,051,772 | 1,243,975 | 2,758 | 36 | 36 |
| 20x |  | 99.88 | 61,358 | 373,335 | 32 | 1 | 0 |
| 2x | 21 | 80.76 | 987,257 | 4,545,450 | 2,924 | 260 | 224 |
|  | 31 | 76.25 | 1,208,314 | 2,823,743 | 4,489 | 379 | 352 |
|  | 41 | 70.78 | 1,478,524 | 2,251,762 | 6,460 | 480 | 447 |
|  | 51 | 64.09 | 1,808,896 | 2,093,319 | 9,115 | 578 | 532 |
|  | 61 | 55.93 | 2,213,535 | 2,230,578 | 12,838 | 709 | 645 |
|  | 71 | 45.94 | 2,710,240 | 2,582,737 | 17,635 | 708 | 628 |

An unsafe unitig is not necessarily a mis-assembly, as it may be a substring of the unsequenced genome by luck. We define an unitig to be *mis-assembled* if its spelling is not a substring of the reference. Table 1 shows that the number of mis-assembled unitigs is substantially lower than the unsafe unitigs, e.g. with 708 mis-assembled unitigs at 2x coverage and *k* = 71. Thus the potential for mis-assembly does not usually translate into a real mis-assembly, though many mis-assemblies remain.

We further check how many of these mis-assembled unitigs fit the example in Fig. 3. A formal definition to capture this example is included in Appendix A for reference. Table 1 shows that the vast majority of mis-assembled cases are in fact caused by this situation, where a repeat has an occurrence in which its start is unsequenced and another occurrence in which its end is unsequenced.

The simulations in Table 1 suggest that the mis-assembly artifact can be removed by simply increasing coverage. In a metagenome expirement, however, this is not always possible. Even when one increases the number of reads, there will continue to be genomes in the sample whose abundance is low enough that their coverage is low. To verify this intuition, we used a standard benchmark dataset generated by the CAMI competition [31], containing 70 million synthetic reads from 30 genomes. Table 2 shows there are 33-37 mis-assembled unitigs, indicating that this artifact remains under realistic coverage of a metagenomic dataset. The section “CAMI dataset” in Appendix C contains more details about the experiment, including Table S1 which shows the details of the dataset.

**Table 2:** The presence of unsafe and mis-assembled unitigs in the CAMI dataset, using simulated error-free reads.

| $k$ | % $k$-mers sampled | n. sequenced segments | n. unitigs | n. unsafe | n. mis-assembled |
| --- | --- | --- | --- | --- | --- |
| 55 | 65 | 171,682 | 174,954 | 593 | 37 |
| 75 | 61 | 207,429 | 207,402 | 656 | 33 |

### Presence of unsafe unitigs in the contig output of real assemblers

We investigated the extent to which the artifact predicted by Theorem 1 appears in output of real assemblers. Assemblers do not simply output the unitigs of a graph but perform many other steps, hence it was not clear if this artifact would appear in the output contigs. Unfortunately, it is not clear how to verify this artifact with real data, as sequencing errors make it difficult to know which of the misassembled contigs are caused by the conditions of Theorem 1. We therefore again used a simulated error-free dataset from the T2T chromosome 1, using the ART simulator [15], with read length of 250 and varying coverages. This time, we simulated reads from either strand, since assemblers are not typically run in single-stranded mode. We also used the CAMI dataset, but simulating reads in double-stranded mode. We then constructed the doubled de Bruijn graph using *k* = 74 and output its maximal unitigs (note that Theorem 1 holds for even *k*). We also ran SPAdes [6] and MEGAHIT [18] to assemble the reads (see Appendix C for parameter details). We then identified unitigs and the assembler contigs that were mis-assembled, but allowing for reverse complements. We will say that a string *x matches* a string *y* with a threshold of *t* if a fraction *t* of the *k*-mers of *x* occur in *y*.

Tables 3 and 4 show that nearly all of the mis-assembled unitigs matched at least one misassembled SPAdes contig with a threshold of 1. For MEGAHIT, the threshold of 1 turned out to be stringent; this is not surprising, since assemblers have many steps that may add or remove *k*-mers from the graph; additionally, MEGAHIT varies the value of *k* internally and may therefore join *k*-mers that do not have an overlap of length *k* −1. Using a threshold of 0.5, however, we found that, similarly to SPAdes, most mis-assembled unitigs matched a mis-assembled contig of MEGAHIT. These results indicate that the artifact predicted by Theorem 1 not only appears in unitigs of the raw graph but also in the output of widely used assemblers like SPAdes and MEGAHIT.

**Table 3:** The extent to which mis-assembled unitigs contribute to mis-assembled contigs of real assemblers. Here, *U* is the set of mis-assembled unitigs in *G*_dbl_, *S* is the set of mis-assembled contigs of SPAdes, and *M* is the set of mis-assembled contigs of MEGAHIT. We use *A* ⊏_*t*_ *B* to indicate the subset of *A* that matches at least one element of *B* at a threshold of *t*.

| | $G_{\text{dbl}}$ | SPAdes | | | MEGAHIT | | |
| --- | --- | --- | --- | --- | --- | --- | --- |
| coverage | $\vert U\vert $ | $\vert S\vert $ | $\vert S \sqsubset_1 U\vert $ | $\vert U \sqsubset_1 S\vert $ | $\vert M\vert $ | $\vert M \sqsubset_{0.5} U\vert $ | $\vert U \sqsubset_{0.5} M\vert $ |
| 1x | 234 | 3,366 | 233 | 209 | 3,070 | 111 | 179 |
| 2x | 129 | 4,423 | 128 | 87 | 2,677 | 75 | 119 |
| 3x | 44 | 8,329 | 44 | 40 | 1832 | 21 | 39 |
| 4x | 13 | 7,365 | 13 | 13 | 1240 | 5 | 13 |
| 5x | 5 | 6,526 | 5 | 5 | 986 | 0 | 5 |
| 6x | 1 | 5,795 | 1 | 1 | 832 | 0 | 1 |

**Table 4:** The extent to which mis-assembled unitigs contribute to mis-assembled contigs of real assemblers for the CAMI metagenomic dataset.

| | $G_{\text{dbl}}$ | metaSPAdes | | | MEGAHIT | | |
| --- | --- | --- | --- | --- | --- | --- | --- |
| $k$ | $\vert U\vert $ | $\vert S\vert $ | $\vert S \sqsubset_1 U\vert $ | $\vert U \sqsubset_1 S\vert $ | $\vert M\vert $ | $\vert M \sqsubset_{0.5} U\vert $ | $\vert U \sqsubset_{0.5} M\vert $ |
| 55 | 37 | 384 | 36 | 36 | 255 | 34 | 32 |
| 75 | 33 | 208 | 30 | 30 | 144 | 30 | 26 |

### Presence of palindrome splitting in a real genome

To measure the extent of the “palindrome splitting” artifact predicted by Theorem 2, we let *K* be the set of all constituent *k*-mers in human chromosome 21 (grch38.p13), after excising the Ns. We confirmed the correctness of Theorem 2 by verifying that the spellings of *D*_non-pal_ are equal to the spellings of *B*_no-loop_ and that the spellings of *B*_last-loop_ are equal to the spellings of *D*_pal_ and are the reverse complements of the spellings of *B*_first-loop_. Table 5 shows that the splitting artifact is present but rare, e.g. for *k* = 15, there were 186 palindromic maximal unitigs in *G*_dbl_(*K*) which were split in *G*_bid_(*K*). The artifact becomes rarer with increasing *k* (e.g. for *k* = 43, there were only 3 split palindromes), which is expected since palindrome frequency in real genomes decreases with length.

**Table 5:** Extent of the palindrome splitting artifact predicted by Theorem 2 in chr21.

| $k$ | $\vert D\vert $ | $\vert B\vert $ | $\vert D_{\text{non-pal}}\vert $ | $\vert D_{\text{pal}}\vert $ | $\vert B_{\text{last-loop}}\vert $ | $\vert B_{\text{first-loop}}\vert $ | $\vert B_{\text{no-loop}}\vert $ |
| --- | --- | --- | --- | --- | --- | --- | --- |
| 15 | 1,465,800 | 1,465,986 | 1,465,614 | 186 | 186 | 186 | 1,465,614 |
| 29 | 60,849 | 60,866 | 60,832 | 17 | 17 | 17 | 60,832 |
| 35 | 36,542 | 36,552 | 36,532 | 10 | 10 | 10 | 36,532 |
| 43 | 18,459 | 18,462 | 18,456 | 3 | 3 | 3 | 18,456 |

### Presence of palindrome splitting in real assemblers

Most assembler papers do not contain enough detail to ascertain what kind of de Bruijn graph they use to handle reverse complements nor what modifications, if any, they make to the unitig algorithm used for the final output. Looking at MEGAHIT [18], SPAdes [6], ABySS [36, 17], and minia [12], only the SPAdes paper is unambiguously clear in saying how it handled reverse complements (it used the doubled dBG). Furthermore, since these assemblers implement many heuristics, the splitting artifact may be absent (respectively, present) even if they did (respectively, did not) use bidirected graphs. We therefore tested the behavior of these assemblers by looking for evidence of palindrome splitting in their output, rather than in their technical descriptions.

Since large exact palindromes are uncommon in typical genomes, we created a synthetic genome by modifying a ∼ 7 mil bp long contig from human chromosome 4 (grch38.p13) as follows. We randomly sampled a 1,000bp-long region and replaced the last 500bp by the reverse complement of first 500 bp; we then repeated the sampling process 700,000 times. We then simulated error-free Illumina reads with ART. We used a read length of 100bp so that assemblers will not be able to supplement the dBG with read information in a way that hides the palindrome splitting artifact. We used 10x coverage so that most *k*-mers would be sampled.

First, we find the reference location of each unitig *w* in *D*_pal_. Then, we find all exact alignments of the assembler contigs to the reference. We say that *w* is *fully-covered* if there exists a contig whose alignment spans *w*’s. Otherwise, we say *w* is *split* if one half of *w*’s region does not overlap with any contig alignments while the other half has a contig aligned that ends precisely in the middle of *w* at one end and extends past *w* at the other end. A unitig is *ambiguous* if it does fall into either category. Appendix C contains a more precise definition of these cases.

Table 6 shows that ABySS clearly exhibits the palindrome splitting artifact, with all non-ambiguous unitigs being split and none fully-covered. In fact, this is due to a heuristic that breaks unitigs at any palindromic edges or vertices [1, 2]. The opposite was true for SPAdes and MEGAHIT, with all non-ambiguous unitigs being fully-covered and none split. minia on the other hand exhibited mixed behavior. Of the 417 non-ambiguous cases, 34 were split and 383 were fully-covered. These results indicate that the palindrome splitting artifact of Theorem 2 does persist all the way to the contig output stage in some assemblers. However, this artifact requires the presence of long exact palindromes in the reference, which is uncommon in most genomes.

**Table 6:**
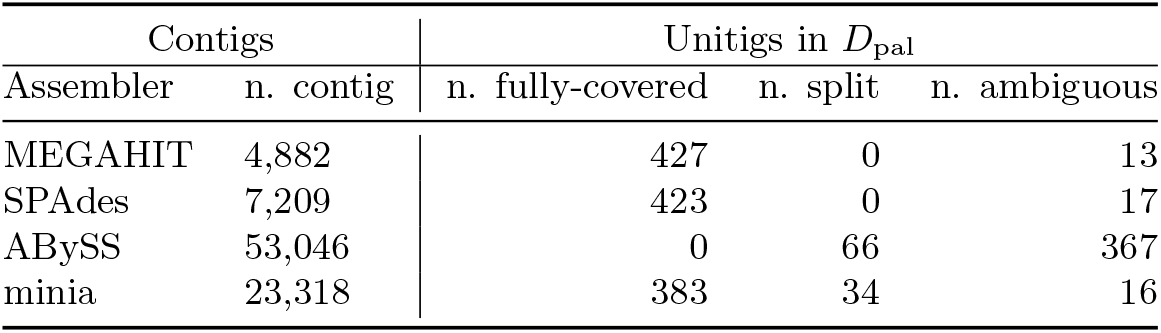
Presence of the palindrome splitting artifact in real assemblers on a synthetic genome. We used *k* = 31 for all the assemblers (see Appendix C for details). We filtered out unitigs shorter than 500 bp, amounting to 440 palindromic strings in ABySS and minia and 433 palindromic strings in SPAdes and MEGAHIT.

## 6 Discussion

Our theoretical study uncovered two artifacts of the unitig algorithm for genome assembly. The first is that even without sequencing errors, it can create mis-assemblies in places of imperfect coverage. The second is that when the bidirected graph is used to model double-strandedness, the unitig algorithm under-assembles by failing to merge the two halves of palindromes. Our experiments confirmed the presence of these theoretically-predicted artifacts in real genomes and popular assemblers. Fortunately, the impact of these artifacts is not large and can be addressed. Mis-assembly issues due to the first artifact can be resolved by increasing coverage or, potentially, breaking unitigs at places where the coverage along them is uneven. Under-assembly issues due to the palindrome artifact are rare in real genomes and, moreover, can sometimes be fixed by forcing the unitigs to “push their way through” lonely inverted loops (however, it is not always possible, e.g. [23, 8].

One of the tangential outcomes of this paper is that we have given proper definitions for things like walks and unitigs in the context of bidirected graphs. Previous papers used these concepts somewhat informally; when definitions were given, they worked in the context of that paper but failed to have more general desired properties. For example, our previous work had an inconsistency in the way that a walk was defined on a single vertex versus on many vertices [28]. One key takeaway is that as a rule thumb, when working with bidirected graphs one should avoid thinking in terms of vertices but think instead of vertex-sides. The definitions we have provided in this paper generalize further than previous ones and are able to form the basis for the type of analysis we have done in this paper. For example, we are the first to prove the bijection between walks in the doubled and bidirected dBGs. We hope that these definitions will facilitate future attempts to formally study questions in bidirected graphs.

Bidirected graphs give an elegant way to capture the double-stranded nature of DNA in a dBG, but our results here indicate that, for the unitig algorithm, they do not give any theoretical advantage. One of the claimed advantages of using the bidirected graph framework in assembly is that it allows one to take advantage of results from graph theory that may otherwise be hidden. The primary example of this is a result (involving one of the authors) in [22] where a variant of the assembly problem was theoretically solved in polynomial time by relying on a reduction to the flow problem in bidirected graphs [14]. When viewed in retrospect, however, it is not clear that this connection was necessary. The algorithm being reduced to [14] was too cumbersome to implement and, when the assembly problem later necessitated a software solution, an approximation algorithm was used instead [20, 21]. But the approximation algorithm worked on the doubled graph, erasing the advantage of having initially formulated the problem on bidirected graphs. Therefore, it remains to be seen if there are situations where the connection to graph theoretical results on bidirected graphs can prove useful for genome assembly. Alternatively, using a different setting may better help identify the advantages of bidirected graphs, e.g. pangenomics [27], rearrangement analysis [7], or compression [29]. Quantifying these advantages would be an exciting future direction.

### Reproducibility

Scripts for the experimental evaluations are available on GitHub [3].

## Acknowledgements

PM thanks Rayan Chikhi, Alexandru Tomescu, and Mihai Pop for useful discussions. This material is based upon work supported by the National Science Foundation under Grant No. 1453527 and 1931531. AR was supported by NIH Computation, Bioinformatics, and Statistics training program.

## A Safety of unitigs: full exposition

### Proof of Theorem 1 and Corollary 1

In this subsection, we will prove Theorem 1 and Corollary 1. In the following, we will always have *𝒮* be a set of sequenced segments and *w* = (*x*_0_, …, *x*_*m*_) be a unitig in *G*_basic_(sp^*k*^(*𝒮*)). We start with a lemma that, roughly speaking, says that if a walk corresponding to some *S* ∈ *𝒮* touches *w*, it must contain all of *w* except that it may begin or end somewhere along the way.

#### Lemma A.1.

*Let 𝒮 be a set of sequenced segments and let w* = (*x*_0_, …, *x*_*m*_) *be a unitig in G*_*basic*_(*sp*^*k*^(*𝒮*)). *Let S* ∈ *𝒮 and let g* = (*g*_0_, …, *g*_|*S*|_) *be the walk corresponding to S. Suppose there exists i and j such that x*_*i*_ = *g*_*j*_. *Then*,

i. *If suf*_*k*_(*S*) ∉ {*x*_*i*_, …, *x*_*m* −1_}, *then g*_*j*+*δ*_ = *x*_*i*+*δ*_ *for all δ* ∈ [0, *m* − *i*].
ii. *If pre*_*k*_(*S*) ∉ {*x*_1_, …, *x*_*i*_}, *then g*_*j* −*δ*_ = *x*_*i* −*δ*_ *for all δ* ∈ [0, *i*].

*Proof*. We will only prove (*i*), since the argument for (*ii*) is symmetric. We use induction on *δ*. For *δ* = 0, we have that the implication of (*i*) reduces to *g*_*j*_ = *x*_*i*_, which is vacuously true because it is also a condition of the theorem. Now we assume that (*i*) holds for *δ* − 1, i.e. *g*_*j*+*δ* −1_ = *x*_*i*+*δ* −1_. Since *x*_*i*+*δ* −1_ ≠ *suf*_*k*_(*G*), *g*_*j*+*δ* −1_ is not the last vertex of *g*. Because *x*_*i*+*δ* −1_ is a non-last vertex of a unitig, it has only one out-neighbor, which is *x*_*i*+*δ*_. Therefore, *g*_*j*+*δ*_ = *x*_*i*+*δ*_, which shows that that (*i*) holds for *δ*. □

Using this lemma, we can now prove some general properties of unsafe unitigs.

#### Lemma A.2.

*Let 𝒮 be a set of sequenced segments and let w* = (*x*_0_, …, *x*_*m*_) *be a unitig in G*_*basic*_(*sp*^*k*^(*𝒮*)). *If w is unsafe then*

i. *m* ≥ 1,
ii. *there exists S* ∈ *𝒮 such that pre*_*k*_(*S*) ∈ {*x*_1_, …, *x*_*m*_},
iii. *there exists S* ∈ *𝒮 such that suf*_*k*_(*S*) ∈ {*x*_0_, …, *x*_*m* −1_},
iv. *for all S* ∈ *𝒮 and their corresponding walks g, either g and w do not share a vertex, or pre*_*k*_(*S*) ∈ {*x*_1_, …, *x*_*m*_}, *or suf*_*k*_(*S*) ∈ {*x*_0_, …, *x*_*m* −1_}, *and*
v. *for all S* ∈ *𝒮 and all i, occ*_*S*_(*x*_*i*_) ≤ 2.

*Proof*. For (*i*), consider a unitig that has just one vertex *x*. Since each *k*-mer in *G*_basic_(sp^*k*^(*𝒮*)), there must be at least one *S* ∈ *𝒮* whose walk contains *x*. Hence, the unitig that is composed of only *x* is safe. For (*ii*), assume for sake of contradiction that for all *S* ∈ *𝒮, pre*_*k*_(*S*) ∉ {*x*_1_, …, *x*_*m*_}. Since every vertex of the graph must be contained in at least one string, let *S*′ ∈ *𝒮* be a string that contains *x*_*m*_. Applying Lemma A.1(*ii*) with *i* = *m*, we get that the walk corresponding to *S*′ must contain *w*, contradicting that *w* is unsafe. The case of (*iii*) is symmetric to (*ii*), using *x*_0_ instead of *x*_*m*_ and applying Lemma A.1(*i*) with *i* = 0. For (*iv*), let *g* = (*g*_0_, …, *g*_|*S*|_) and assume for sake of contradiction that there exists a *S* ∈ *𝒮* such that *g* shares a vertex with *w* and *pre*_*k*_(*S*) ∉ {*x*_1_, …, *x*_*m*_} and *suf*_*k*_(*S*) ∉ {*x*_0_, …, *x*_*m* −1_}. Let *x*_*i*_ and *g*_*j*_ be the vertices of *w* and *g*, respectively, that are equivalent. We can apply Lemma A.1 to get that (*g*_*j*_, …, *g*_*j*+*m* −*i*_) = (*x*_*i*_, …, *x*_*m*_) and (*g*_*j* −*i*_, …, *g*_*j*_) = (*x*_0_, …, *x*_*i*_). This means that *w* is a subwalk of *g*, which is a contradiction. For (*v*), let *S* ∈ *𝒮* and let *g* = (*g*_0_, …, *g*_|*S*|_) be its corresponding walk. If *g* and *w* do not share any vertices, then *occ*_*S*_(*x*_*i*_) = 0 ≤ 2 for all *i* and we are done. Otherwise, we can apply (*iv*) to get that either (1) *pre*_*k*_(*S*) ∈ {*x*_1_, …, *x*_*m*_} or (2) *suf*_*k*_(*S*) ∈ {*x*_0_, …, *x*_*m* −1_}. Let us consider (1) — we will omit the argument for (2) since it is symmetrical. Then *g*_0_ = *x*_*i*_ for some 1 ≤ *i* ≤ *m*. Note that *g*_0_ is the first occurrence of *x*_*i*_ in *g*. Assume for the sake of contradiction that *occ*_*S*_(*x*_*i*_) *>* 2. To get the second occurrence of *x*_*i*_, *g* must first visit *x*_0_. After this second visit to *x*_*i*_, *g* must continue all the way until *x*_*m*_ if it is to visit *x*_*i*_ for a third time. Therefore, at the second visit to *x*_*i*_, *g* must in fact visit (*x*_0_, …, *x*_*m*_), which contradicts that *w* is unsafe.

The case when a sequenced segment contains its first and/or last *k*-mer more than once puts additional constraints on how it can contain a unitig.

#### Lemma A.3.

*Let 𝒮 be a set of sequenced segments and let w* = (*x*_0_, …, *x*_*m*_) *be a unitig in G*_*basic*_(*sp*^*k*^(*𝒮*)). *Let S* ∈ *𝒮 such that at least one of the following holds:*

i. *occ*_*S*_(*pre*_*k*_(*S*)) = 2 *and there exists an integer i* ∈ [1, *m*] *such that x*_*i*_ = *pre*_*k*_(*S*), *or*
ii. *occ*_*S*_(*suf*_*k*_(*S*)) = 2 *and there exists an integer j* ∈ [*i, m* − 1] *such that x*_*j*_ = *suf*_*k*_(*S*). *Then, spell*(*w*) *is not a substring of S iff both* (*i*) *and* (*ii*) *hold*.

*Proof*. We only prove case (*i*) since case (*ii*) is symmetrical. Let *g* be the walk corresponding to *S*. In the first phase, *g* starts from *x*_*i*_ and, since it must visit *x*_*i*_ a second time, continues until *x*_*m*_. Then at some point it enters *w* through *x*_0_ and proceeds to visit *x*_*i*_ for the second and last time. We will refer to the time from the end of the first phase to the point it enters *x*_0_ as the second phase, and the rest of the walk as the third phase. Observe that *g* does not contain *w* as a subwalk in either the first or second phase.

Now we prove the if direction. During phase 1, *g* visits *x*_*j*_ exactly once. During phase 2, *g* does not visit *x*_*j*_. During phase 3, *g* proceeds from *x*_0_ forward along the unitig until it hits *x*_*j*_ for the second time. Since *x*_*j*_ is occurs exactly twice and is the last vertex of *g*, this is the end of *g*. Since *j < m, g* does not contain *w* as a subwalk during the third phase.

Now we prove the only if direction. Assume *w* is not a subwalk of *g*. Therefore, during the third phase *g* cannot go until *x*_*m*_ and must stop earlier at some *x*_*j*_ = *suf*_*k*_(*S*), for some integer *j* ∈ [*i, m* − 1]. This *x*_*j*_ was visited once during phase 1 and not visited during phase 2 and now visited a second and final time during phase 3. □

These lemmas are all the pieces we need to prove Theorem 1.

#### Theorem 1.

*Proof*. First we prove the if direction. We will show that for all *S* ∈ *𝒮* and its corresponding walk *g*, if one of the four conditions hold, then *w* is not a subwalk of *g*. If (*i*) holds, then *w* is trivially not a subwalk of *g*. Now, if *pre*_*k*_(*S*) = *x*_*i*_ for some 1 ≤ *i* ≤ *m* and *x*_*i*_ is visited only once by *g*, If (*ii*) holds, then *g* starts with *x*_*i*_ but never visits *x*_*i*_ again, therefore (*x*_0_, …, *x*_*i*_) is not a subwalk of *g*. Hence, *w* is not a subwalk of *g*. Similarly, if (*iii*) holds, then (*x*_*j*_, …, *x*_*m*_) is not a subwalk of *g* and hence *w* is not a subwalk of *g*. If (*iv*) holds, then Lemma A.3 implies that *w* is not a subwalk of *g*.

Now we prove the only if direction. We will show that for all *S* ∈ *𝒮* and their corresponding walk *g*, if *w* is not a subwalk of *g*, then one of the four conclusions hold. By Lemma A.2.(*iv*), either (1) *g* does not contain any *k*-mer from *w*, (2) *pre*_*k*_(*S*) ∈ {*x*_1_, …, *x*_*m*_}, or (3) *suf*_*k*_(*S*) ∈ {*x*_0_, …, *x*_*m* −1_}. In case of (1), *g* trivially does not contain *w*, and condition (*i*) is satisfied. In case of (2), let *i* ∈ [1, *m*] be an integer such that *x*_*i*_ = *pre*_*k*_(*S*). By Lemma A.2.(*v*), *occ*_*S*_(*x*_*i*_) is either 1 or 2. If *occ*_*S*_(*x*_*i*_) = 1, then condition (*ii*) immediately holds. If *occ*_*S*_(*x*_*i*_) = 2, then Lemma A.3 implies that there exists an integer *j* ∈ [*i, m* − 1] that satisfies condition (*iv*). In case of (3), let *j* ∈ [0, *m* − 1] be an integer such that *x*_*j*_ = *suf*_*k*_(*S*). Again, by Lemma A.2.(*v*), *occ*_*S*_(*x*_*j*_) is either 1 or 2. If *occ*_*S*_(*x*_*j*_) = 1, then condition (*iii*) immediately holds. If *occ*_*S*_(*x*_*j*_) = 2, then Lemma A.3 implies that there exists an integer *i* ∈ [*i, m* − 1] that satisfies condition (*iv*). □

#### Corollary 1.

*Proof*. We can apply Theorem 1 to set *𝒮* = {*X*}. Since *X* must contain all *k*-mers of *w, w* is unsafe if and only if condition (ii), (iii) or (iv) from Theorem 1 holds for *S* = *X*. First, assume Condition (ii) is true for *X*. Then by Theorem 1, *w* is unsafe. Consider the walk *g* corresponding to *X*. Because *g* begins at *x*_*i*_ and all vertices in *G*_basic_(*K*) must be in *g* at least once, (*x*_*i*_, …, *x*_*m*_) is a subwalk of *g*. This is the one and only occurrence of *x*_*i*_ in *g*. Since *x*_*i*_ is the first vertex in *g* occuring only once, *x*_*i* −1_ cannot precede *x*_*i*_. Hence, *x*_*i* −1_ must be the end of *g*, i.e., *suf*_*k*_(*S*) = *x*_*i* −1_. Note that, this is the one and only occurrence of *x*_*i* −1_ in *g*. Thus, Condition (iii) is also true for *X*. With a symmetric argument, we can show that if Condition (iii) is satisfied, then Condition (ii) is satisfied. Combining both gives us the first condition of the corollary. Finally, observe that Condition (iv) is identical to Condition 2 in the corollary. The fact that these conditions can hold for at most one unitig follows directly from the fact that there is only one vertex for *pre*_*k*_(*S*) in the graph.

### Formal definition of the case of Figure 3

In Section 5, we quantify the number of unsafe unitigs that fall into the case of Figure 3. To make this precise, we give a formal classification for this case. Let *X* be a genome and let *𝒮* be a set of its sequenced segments. We say that an unsafe walk *w* = (*x*_0_, …, *x*_*m*_) satisfies the case of Figure 3 if

i. there exists 0 *< i* ≤ *j < m* such that ψ = (*x*_*i*_, …, *x*_*j*_) is a unitig in *G*_basic_(sp^*k*^(*X*)),
ii. *spell*(ψ) occurs at least twice in *X*,
iii. in one of the occurrences, the *k*-mer preceding *spell*(ψ) is not in *𝒮*,
iv. in another of the occurrences, the *k*-mer following *spell*(ψ) is not in *𝒮*,
v. there exists *S* ∈ *𝒮* and an integer *i*′ ∈ [*i, j*] such that *spell*((*x*_*i*′_, …, *x*_*j*_)) is a suffix of *S*, and
vi. there exists *S* ∈ *𝒮* and an integer *j*′ ∈ [*i*′ − 1, *j*] such that *spell*((*x*_*i*_, …, *x*_*j*′_)) is a suffix of *S*.

**Figure S1:**
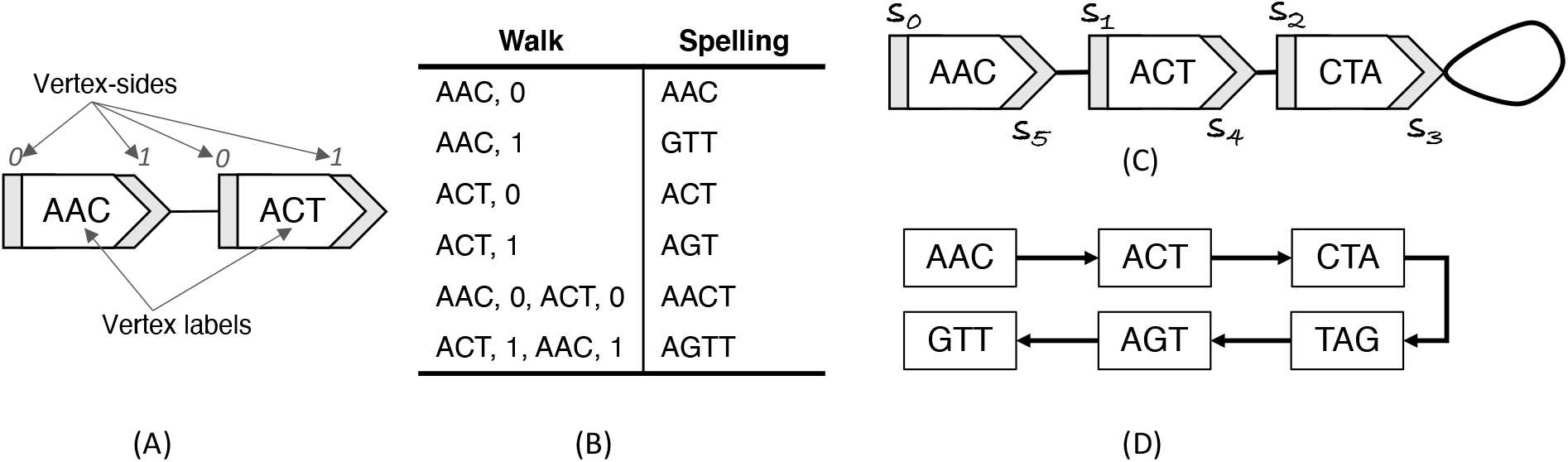
An example illustrating some of the bidirected graph terminology. Panel (A) shows a bidirected graph with two vertices. Panel (B) shows a list of all possible walks in this graph and their spellings. Note that walk (*AAC*, 0, *ACT*, 0) and (*ACT*, 1, *AAC*, 1) are reverses of each other. The endpoint sides of the walks, in both cases, are (*AAC*, 0) and (*ACT*, 1). Panel C and D show an example of a doubled dBG (*G*_dbl_(*K*)) and a bidirected dBG (*G*_bid_(*K*)) using *K* = {*AAC, ACT, CT A*}. Panel C shows *G*_bid_(*K*) as well as the order of vertex-sides as they appear in a walk *w* = (*AAC*, 0, *ACT*, 0, *CT A*, 0, *CT A*, 1, *ACT*, 1, *AAC*, 1) = (*s*_0_, …, *s*_5_), with *spell*(*w*) = *AACT AGTT*. Note that in this case, since *spell*(*w*) is a palindrome, the reverse walk is identical: *rev*(*w*) = *w*. Panel D shows *G*_dbl_(*K*).

## B The relationship of *G*_dbl_(*K*) and *G*_bid_(*K*): full exposition

In this section, we will prove Theorem 2. We start by providing additional definitions that are necessary to understand the proofs in this section.

Let *K* be a set of *k*-mers. A unitig in a directed graph that is not a proper subwalk of another unitig that ends at the same vertex is said to be *prefix-maximal*; a unitig that is not a proper subwalk of another unitig that starts with the same vertex is said to be *suffix-maximal*. Notice that a unitig is maximal iff it is both prefix- and suffix-maximal.

Let (*u, s*) be a vertex-side in *G*_bid_(*K*). We define *d*^*il*^(*u, s*) to indicate the presence of an inverted loop, i.e. *d*^*il*^(*u, s*) = 1 if there is an inverted loop incident to side (*u, s*) and *d*^*il*^(*u, s*) = 0 otherwise. A unitig *t* in *G*_bid_(*K*) is *prefix-maximal* if it is not a proper subwalk of another unitig that ends at the same vertex-side as *t*. A unitig is *suffix-maximal* if it is not a proper subwalk of another unitig that starts with the same vertex-side as *t*. Note that a unitig is maximal iff it is both prefix- and suffix-maximal.

We will prove Theorem 2 by first building a collection of Lemmas. First, we make a simple observation. A palindrome must have an even number of characters, otherwise there is a middle character that would need to be equal to its own reverse complement. Hence, a palindromic walk, in either the doubled or the bidirected graph must have an even number of nucleotides.

### Lemma B.1.

*Let K be a set of k-mers*.

1. *For all palindromic walks w* = (*x*_0_, …, *x*_*n*_) *in G*_*dbl*_(*K*), *k and n have the same parity*.
2. *For all palindromic walks t* = (*u*_0_, *s*_0_, …, *u*_*n*_, *s*_*n*_) *in G*_*bid*_(*K*), *k and n have the same property*.

*Proof*. A palindromic string must have an even number of nucleotides. The number of nucleotides in *spell*(*w*) and in *spell*(*t*) is *k* + *n*, Hence the parity of *k* and *n* must be the same.

From here on out, we proceed by first proving Lemmas for the directed de Bruijn graphs (both the regular one and the doubled one) (Appendix B.1), then proving Lemmas for the bidirected graph (Appendix B.2), then proving Lemmas which connect the two types of graphs (Appendix B.3), and, finally, proving Theorem 2 (Appendix B.4).

### B.1 Directed graph

First, we make the observation that unitigs cannot repeat vertices unless they are a simple cycle. This is generally stated without proof, but the statement is actually not true when unitigs are allowed to be periodic cycles. In our definition of unitig, we forbid this case, allowing us to prove the observation.

#### Lemma B.2.

*For all unitigs w in a directed graph, either w is a simple cycle or it does not repeat any vertices*.

*Proof*. Let *w* = (*x*_0_, …, *x*_*n*_) be a unitig. Suppose that *w* repeats a vertex. Let 0 ≤ *j* ≤ *n* be the smallest value for which there exists 0 ≤ *i < j* such that *x*_*i*_ = *x*_*j*_. If *i >* 0, then *x*_*i*_ has *x*_*i* −1_ and *x*_*j* −1_ as an in-neighbor. By the minimality of our choice of *i, x*_*i* −1_≠ *x*_*j* −1_, and hence *d*^−^(*i*) ≥ 2. This contradicts that *w* is a unitig. If *i* = 0, then let *j* + 1 ≤ 𝓁 ≤ *n* − 1 be the largest index greater than *j* such that *x*_𝓁_ = *x*_𝓁 mod (*j*+1)_. In other words, 𝓁 is the first place after *x*_*j*_ where the unitig is about to “fall off the cycle”. If such an 𝓁 does not exist, then either *j* = *n* and *w* is a simple cycle, or *w* is a simple periodic cycle, contradicting the definition of a unitig. Otherwise, the vertex *x*_𝓁_ has as out-neighbors both *x*_𝓁+1_ and *x*_𝓁+1 mod (*j*+1)_. By the choice of 𝓁, these out-neighbors are distinct and hence *d*^+^(*x*_𝓁_) ≥ 2. This contradicts that *w* is a unitig.

A very simple property in the doubled graph is is that the in-degree (respectively, out-degree) of a vertex is equal to the out-degree (respectively, in-degree) of its reverse complement.

#### Lemma B.3.

*Let K be a set of k-mers and let x be a vertex in G*_*dbl*_(*K*). *Then d*^+^(*x*) = *d*^−^(*x*) *and d*^−^(*x*) = *d*^+^(*x*).

*Proof*. Observe that for all vertices *y* in the *G*_dbl_(*K*), there is an edge from *x* to *y* in *G*_dbl_(*K*) iff there is an edge from 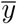 to 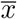. This is true even if 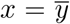 and these two edges are identical. Hence 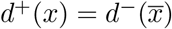 and 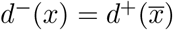. □

We defined maximal unitigs as those that are not proper sub-walks of other unitigs. We can give an equivalent definition for directed graphs, in terms of vertex degrees. Since it is widely known, we state it without proof.

#### Lemma B.4.

*Let G be a directed graph and let w* = (*x*_0_, …, *x*_*n*_) *be a unitig in G. Then*

i. *w is prefix-maximal if and only if d*^−^(*x*_0_) ≠ 1 *or there exists a vertex x*′ *that has an edge to x*_0_ *and d*^+^(*x*′) *>* 1.
ii. *w is suffix-maximal if and only if d*^+^(*x*_*n*_) ≠ 1 *or there exists a vertex x*′ *that has an edge from x*_*n*_ *and d*^−^(*x*′) *>* 1.

Palindromic unitigs play a special role in Theorem 2. We observe that in a palindromic unitig of the doubled graph, the only edge from a *k*-mer to its reverse complement is the middle one.

#### Lemma B.5.

*Let K be a set of k-mers with odd k. Let w* = (*x*_0_, …, *x*_*n*_) *be a palindromic unitig in G*_*dbl*_(*K*) *that is not a simple cycle. Then for all* 0 ≤ *i* ≤ *n* − 1, *we have that* 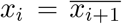 *iff i* = (*n* − 1)*/*2.

*Proof*. First note that by Lemma B.1, *n* is odd and *n* ≥ 1. Let *m* = (*n* − 1)*/*2. Because *spell*(*w*) is a palindrome, 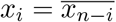 for all 0 ≤ *i* ≤ *n*. The only if direction of the Lemma statement follows immediately by plugging in *i* = *m* and getting 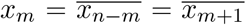. For the if direction, assume that 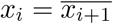 for all 0 ≤ *i* ≤ *n* − 1. Then 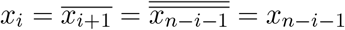. By the fact that *w* is not a simple cycle and Lemma B.2, it cannot have any repeated vertices. Hence, *i* = *n* − *i* − 1 which only happens when *i* = *m*. □

We also observe that a maximal unitig that is not a palindrome cannot contain within it a palindrome of length ≥ *k*.

#### Lemma B.6.

*A non-palindromic maximal unitig w in G*_*dbl*_(*K*) *cannot contain a proper sub-unitig that is palindromic*.

*Proof*. For the sake of contradiction, let *z* be a proper sub-unitig of *w* that is a palindrome. First suppose that there exists a *k*-mer *y* such that *y* precedes *z* in *w* and 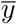 follows *z* in *w*. In that case, observe that the walk 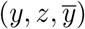 is also a sub-unitig of *w* and also a palindrome. We can then extend *z* in this way until no longer possible, i.e. there do not exist a *k*-mer *y* such that *y* precedes *z* in *w* and 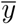 follows *z* in *w*. Let *w*′ be this maximally extended walk. Note that by construction, *w*′ is a sub-unitig of *w* and it is proper because *w*′ is palindromic and *w* is not. Let the first vertex of *w*′ be *x*, and, hence, the last one is 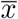.

Consider the case when *w* starts with *x*. Because *w*≠ *w*′, there must exist an out-neighbor *u* of 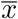 in *w*. Its mirror must also exist, i.e. an edge from *ū* to *x*. Lemma B.4 states that *x* is the first vertex of a maximal unitig, it must either (a) have one other in-neighbor besides *ū* or (b) *ū* must have at least one other out-neighbor besides *x*. For case (a), Lemma B.3 implies that 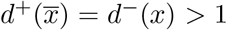. For case (b), Lemma B.3 implies that 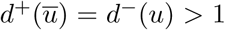. In either case, the degrees of *x* or of *u* contradict the definition of being part of a unitig. The case when *w* ends with 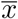 is symmetric and omitted.

Now consider the case when *w* does not start with *x* and does not end with 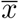. Let *a* be the vertex preceding *x* in *w*, and let *b* be the vertex following 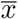 in *w*. There exist a mirror edge from 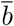 to *x*. Since *w*′ was chosen so that it cannot be extended, *ā* ≠ *b*. Hence *x* has two distinct in-neighbors, *a* and 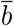. Since *w* contains *x* as a non-first vertex, this contradicts that *w* is a unitig. □

### B.2 Bidirected graph

As is the case with directed graphs (Lemma B.4), there is a definition of maximality for bidirected unitigs that has to do with degrees rather than sub-unitigs. We are not aware of this equivalence being explicitly proven, so we do so here:

#### Lemma B.7.

*Let K be a set of canonical k-mers. Let t* = (*u*_0_, *s*_0_, …, *u*_*n*_, *s*_*n*_) *be a unitig in G*_*bid*_(*K*). *Then*

i. *t is prefix-maximal if and only if d*(*u*_0_, *s*_0_) ≠ 1 *or there is an edge* {(*u*_0_, *s*_0_), (*u*′, *s*′)} *such that d*(*u*′, *s*′) *>* 1, *and*
ii. *t is suffix-maximal if and only if d*(*u*_*n*_, 1 − *s*_*n*_)≠ 1 *or there is an edge* {(*u*_*n*_, 1 − *s*_*n*_), (*u*′, *s*′)} *such that d*(*u*′, *s*′) *>* 1.

*Proof*. We will only prove (*i*) since the proof of (*ii*) is symmetric. First, we prove the only if direction. We need to consider three cases. The first case is when *d*(*u*_0_, *s*_0_)≠ 1. If *d*(*u*_0_, *s*_0_) = 0, then *t* is prefix-maximal because there is no other walk of which it is a subwalk with the same last vertex-side (*u*_*n*_, *s*_*n*_). The second case is when *d*(*u*_0_, *s*_0_) *>* 1. Consider any walk *t*′ that ends in (*u*_*n*_, *s*_*n*_) and of which *t* is a proper subwalk. Observe that (*u*_0_, *s*_0_) would not be the first vertex-side of *t*′. Therefore, since *d*(*u*_0_, *s*_0_) *>* 1, *t*′ cannot be a unitig and *t* must be prefix-maximal. The third case is when (*u*_0_, *s*_0_) has degree one and (*u*′, *s*′) is its only neighbor. Again, consider any walk *t*′ that ends in (*u*_*n*_, *s*_*n*_) and of which *t* is a proper subwalk. Observe that (*u*′, 1 − *s*′) belongs to *t*′ but is not the last vertex-side of *t*′. Therefore, since we assumed that *d*(*u*′, *s*′) *>* 1, *t*′ cannot be a unitig and *t* must be prefix-maximal.

To prove the if direction we prove the contrapositive. In other words, we will show that if the degree of (*u*_0_, *s*_0_) is one and its sole neighbor (*u*′, *s*′) also has degree at most 1, then *t* is not prefix-maximal. First, observe that *t*′ = (*u*′, 1 − *s*′, *u*_0_, *s*_0_, …, *u*_*n*_, *s*_*n*_) is a valid walk, since the edge {(*u*′, *s*′), (*u*_0_, *s*_0_)} exists. Then, observe that the degree of (*u*′, *s*′) is exactly one because it has degree at most one (by our assumption) and also has a neighbor (i.e. (*u*_0_, *s*_0_)). Therefore, the degree requirements for *t*′ being a unitig are fulfilled. Finally, observe that *t* is a proper subwalk of *t*′ ending in the same vertex-side, (*u*_*n*_, *s*_*n*_). Therefore, *t* is not prefix-maximal. □

In a bidirected graph, a walk and its reverse are either both unitigs or not and, if they are, are either both are maximal or not.

#### Lemma B.8.

*Let K be a set of canonical k-mers and let w be a unitig in G*_*bid*_(*K*).

i. *rev*(*w*) *is a unitig in G*_*bid*_(*K*).
ii. *w is prefix-maximal iff rev*(*w*) *is suffix-maximal*.
iii. *w is suffix-maximal iff rev*(*w*) *is prefix-maximal*.

*Proof*. Let (*u*_0_, *s*_0_, …, *u*_*n*_, *s*_*n*_) = *w* and 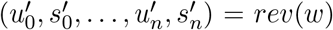. For (*i*), by definition of *rev*, we have that 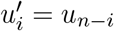 and 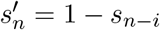. Applying the definition of unitig to *w*, we get that

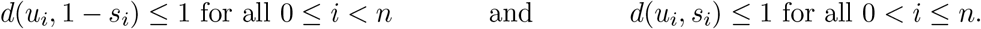

These can be equivalently stated as

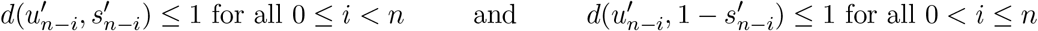

If we change the index variables, these can be equivalently restated as

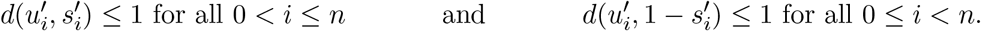

This is precisely the definition of *rev*(*w*) being a unitig.

For (*ii*) and (*iii*), first observe that Lemma B.7 gives an alternate, equivalent, definition for prefix- and suffix-maximal. For (*ii*), observe that if apply the alternate definition of suffix-maximal to *rev*(*w*) and plug in that 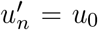 and 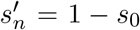, we get precisely the alternate definition of *w* being prefix-maximal. For (*iii*), observe that if apply the alternate definition of prefix-maximal to *rev*(*w*) and plug in that 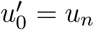_*n*_ and 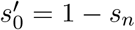, we get precisely the alternate definition of *w* being suffix-maximal.

While we showed that it is natural for the doubled graph to have a palindromic unitig, this is impossible in a bidirected graph.

#### Lemma B.9.

*Let K be a set of canonical k-mers, with k odd. Then a unitig of G*_*bid*_(*K*) *cannot be a palindrome*.

*Proof*. Let *t* = (*u*_0_, *s*_0_, …, *u*_*n*_, *s*_*n*_) be a palindromic walk. By Lemma B.1, *n* is odd, and so *n* ≥ 1. For convenience, let *m* = (*n* − 1)*/*2. By definition, 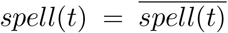. In particular, the two “central” *k*-mers of *spell*(*t*) must be reverse complements of each other. Formally, *orient*(*lab*(*u*_*m*_), *s*_*m*_) = *orient*(*lab*(*u*_*m*+1_), *s*_*m*+1_). Since the labels of vertices in a bidirected graph are distinct, *lab*(*u*_*m*_) ≠ *lab*(*u*_*m*+1_) and hence *s*_*m*_ = 1 − *s*_*m*+1_. Applying the definition of a bidirected walk to *t*, we get that {(*u*_*m*_, 1 − *s*_*m*_), (*u*_*m*+1_, *s*_*m*+1_)} is an edge. The fact that *s*_*m*_ = 1 − *s*_*m*+1_ implies that this edge is an inverted loop incident to (*u*_*m*_, 1 − *s*_*m*_). Thus *d*(*u*_*m*_, 1 − *s*_*m*_) ≥ 2, implying that *t* does not satisfy the definition of being a unitig. □

### B.3 Connecting the directed and bidirected graphs

So far, we have proven properties of the doubled graph and of the bidirected graph separately; in this section, we prove lemmas about the relationship between the two graphs, when *k* is odd. Recall that for a *k*-mer *x* ∈ *K*, we defined *F*_*V*_ (*x*) = (*u, s*), where (*u, s*) is the unique vertex-side in *G*_bid_(*K*) such that *lab*(*u*) = *orient*(*x, s*).

#### Lemma B.10.

*Let K be a set of canonical k-mers where k is odd. F*_*V*_ *is a bijection between vertices of G*_*dbl*_(*K*) *and vertex-sides of G*_*bid*_(*K*).

*Proof*. To show that *F*_*V*_ is a bijection, we will show that for all vertex-sides (*u, s*) in *G*_bid_(*K*), there exists a unique *k*-mer *x* in *G*_dbl_(*K*) such that *F*_*V*_ (*x*) = (*u, s*). Consider a value of *x* such that *F*_*V*_ (*x*) = (*u, s*). By definition, *lab*(*u*) = *orient*(*x, s*)). Since *k* is odd and *x* is not a palindrome, the value of *x* satisfying this must be unique. By construction of *G*_dbl_(*K*) and *G*_bid_(*K*), *k* must be a vertex in *G*_dbl_(*K*). Further, if *x* = *orient*(*lab*(*u*), *s*), then *orient*(*x, s*) = *orient*(*orient*(*lab*(*u*), *s*), *s*) = *lab*(*u*) and so *x* satisfies the condition that *F*_*V*_ (*x*) = (*u, s*).

We will use 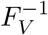 to denote the inverse of *F*_*V*_, which was shown in Lemma B.10 is 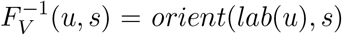. We will use 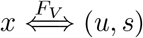 to denote that a vertex *x* of *G*_dbl_ (*K*) and a vertex-side (*u, s*) in *G*_bid_(*K*) are associated with each other by *F*_*V*_.

Recall that for two *G*_dbl_(*K*) *k*-mers *x*_1_ and *x*_2_, we define the mapping *F*_*E*_(*x*_1_, *x*_2_) = {(*u*_1_, 1 − *s*_1_), (*u*_2_, *s*_2_), where (*u*_1_, *s*_1_) = *F*_*V*_ (*x*_1_) and (*u*_2_, *s*_2_) = *F*_*V*_ (*x*_2_). Though the mapping is not a bijection, it preserves the property of being an edge in the respective graph^2^:

#### Lemma B.11.

*Let K be a set of canonical k-mers where k is odd. Let x*_1_ *and x*_2_ *be vertices in G*_*dbl*_(*K*). *We have that* (*x*_1_, *x*_2_) *is an edge in G*_*dbl*_(*K*) *if and only if F*_*E*_(*x*_1_, *x*_2_) *is an edge in G*_*bid*_(*K*).

*Proof*. By the definition of bidirected edges, *F*_*E*_(*x*_1_, *x*_2_) = {(*u*_1_, 1 − *s*_1_), (*u*_2_, *s*_2_)} is an edge iff

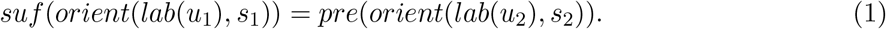

Recall that by the definition of *F*_*E*_, *lab*(*u*_1_) = *orient*(*x*_1_, *s*_1_) and *lab*(*u*_2_) = *orient*(*x*_2_, *s*_2_). We can therefore rewrite Equation (1) equivalently as

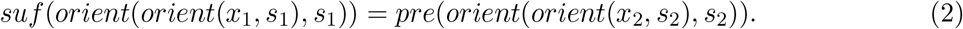

Now, using the fact that *orient*(*orient*(*y, s*), *s*) = *y*, for all *y* and *s*, we can rewrite Equation (2) as

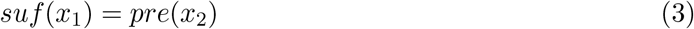

Since we obtained Equation (3) from Equation (1) using equivalent transformations, it shows that the two statements are equivalent and completes the proof. □

One particular case of Lemma B.11 that we will often invoke is that there is an edge from *x* to 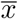 in *G*_dbl_(*K*) if and only if there is an inverted loop incident to (*u*, 1 − *s*) in *G*_bid_(*K*).

Now recall that *F*_*W*_ is defined as a function that maps a walk *w* = (*x*_0_, …, *x*_*n*_) in *G*_dbl_(*K*) to a sequence *F*_*W*_ (*w*) = (*u*_0_, *s*_0_, …, *u*_*n*_, *s*_*n*_), with (*u*_*i*_, *s*_*i*_) = *F*_*V*_ (*x*_*i*_) for all 0 ≤ *i* ≤ *n*. We show that *F*_*W*_ (*w*) is in fact a walk in *G*_bid_(*K*) and, moreover, *F*_*W*_ is a bijection from the set of walks in *G*_dbl_(*K*) to the set of walks in *G*_bid_(*K*).

#### Lemma B.12.

*Let K be a set of canonical k-mers where k is odd. F*_*W*_ *is a spell-preserving bijection from the set of walks in G*_*dbl*_(*K*) *to the set of walks in G*_*bid*_(*K*).

*Proof*. Let *w* = (*x*_0_, …, *x*_*n*_) be a walk in *G*_dbl_(*K*) and let (*u*_*i*_, *s*_*i*_) = *F*_*V*_ (*x*_*i*_) for all 0 ≤ *i* ≤ *n*. We will first show that *F*_*W*_ (*w*) = (*u*_0_, *s*_0_, …, *u*_*n*_, *s*_*n*_) is a walk in *G*_bid_(*K*). By definition of *F*_*V*_, *F*_*W*_ (*w*) is a sequence of vertex-sides. Consider the edge from *x*_*i*_ to *x*_*i* −1_, for all 1 ≤ *i* ≤ *n*. By Lemma B.11, there is an edge {(*u*_*i* −1_, 1 −*s*_*i* −1_), (*u*_*i*_, *s*_*i*_)} in *G*_bid_(*K*). This shows that every two consecutive vertex-sides in *F*_*W*_ (*w*) are connected by an edge, thus completing the proof that *F*_*W*_ (*w*) is a walk. The fact that it is spell preserving follows from its definition.

To show that *F*_*W*_ is a bijection, we need to show that for all walks *t* = (*u*_0_, *s*_0_, …, *u*_*n*_, *s*_*n*_) in *G*_bid_(*K*), there exists a unique walk *w* in *G*_dbl_(*K*) such that *t* = *F*_*W*_ (*w*). Let *w* = (*x*_0_, …, *x*_*n*_) be an arbitrary walk in *G*_dbl_(*K*). In order for *F*_*W*_ (*w*) = *t*, we need that *F*_*V*_ (*x*_*i*_) = (*u*_*i*_, *s*_*i*_) for all 0 ≤ *i* ≤ *n*. Because *F*_*V*_ is bijection (Lemma B.10), there is exactly one value of *x*_*i*_ to satisfy this, and that is *x*_*i*_ = *F* ^−1^(*u*_*i*_, *s*_*i*_) = *orient*(*lab*(*u*_*i*_), *s*_*i*_). Therefore, *w* = (*orient*(*lab*(*u*_0_), *s*_0_), …, *orient*(*lab*(*u*_*n*_), *s*_*n*_)) is the unique walk in *G*_dbl_(*K*) to satisfy *F*_*W*_ (*w*) = *t*. □

Given the above proof, we can write the inverse of *F*_*W*_ as 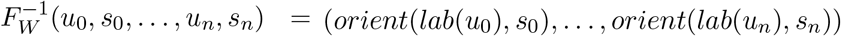. We will use 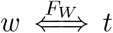 to denote that a walk *w* in *G*_dbl_(*K*) and a walk *t* in *G*_bid_(*K*) are associated with each other by *F*_*W*_.

Notice that if *k* were to be even, then Lemma B.12 would not hold. In particular, Let *x* ∈ *K* be a palindrome *k*-mer and let *u* be the vertex in *G*_bid_(*K*) such that *lab*(*u*) = *x*. Then both of the walks (*u*, 0) and (*u*, 1) would spell *x*, while in the *G*_dbl_(*K*) there would only be one walk that spells *x*.

Since unitigs are defined in terms of degrees, it is useful to first understand how the degrees of vertices in *G*_dbl_(*K*) relate to the degrees of vertex sides in *G*_bid_(*K*).

#### Lemma B.13.

*Let K be a set of canonical k-mers where k is odd. Let x be a vertex in G*_*dbl*_(*K*) *and let* (*u, s*) *be a vertex-side is G*_*bid*_ (*K*) *such that* 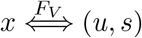. *Then*,

i. *d*^+^(*x*) = *d*(*u*, 1 − *s*) − *d*^*il*^(*u*, 1 − *s*)
ii. *d*^−^(*x*) = *d*(*u, s*) − *d*^*il*^(*u, s*).

*Proof*. For proving part (*i*), we will first prove an upper bound and then a matching lower bound. We start with the upper bound. Let *Y* be the set of all out-neighbors of *x* which are not equal to 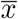. Note that *Y* may contain *x*. Let *Y*′ = {*F*_*V*_ (*y*) | *y* ∈ *Y*} and observe that since *F*_*V*_ is injective (Lemma B.10), |*Y*′| = |*Y* |. By Lemma B.11, for each vertex-side (*u*′, *s*′) ∈ *Y*′, there is an edge {(*u*, 1 − *s*), (*u*′, *s*′)} and so *d*(*u*, 1 − *s*) ≥ |*Y*′|.

We show that *d*^+^(*x*) = *d*(*u*, 1 − *s*) − *d*^*il*^(*u*, 1 − *s*) by considering two cases. In the first case, assume that there does not exist an edge 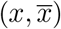. Then *d*^+^(*x*) = |*Y* |. Moreover, by Lemma B.11, the edge {(*u*, 1 − *s*), (*u*, 1 − *s*)} does not exist, so *d*^*il*^(*u*, 1 − *s*) = 0. Putting these facts together, *d*^+^(*x*) = |*Y* | = |*Y*′| ≤ *d*(*u*, 1 − *s*) = *d*(*u*, 1 − *s*) − *d*^*il*^(*u*, 1 − *s*).

In the second case, assume that there exists an edge 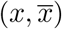. Lemma B.11 says that there is an inverted loop incident to side (*u*, 1 − *s*), so *d*^*il*^(*u*, 1 − *s*) = 1. An inverted loop adds 2 to the degree of (*u*, 1 − *s*), i.e. *d*(*u*, 1 − *s*) ≥ |*Y*′| + 2; it also contributes 1 to out-degree of *x*, i.e. *d*^+^(*x*) = |*Y* | + 1. Putting these together, we get *d*^(^*x*) = |*Y* | + 1 = |*Y*′| + 1 ≤ *d*(*u*, 1 − *s*) − 1 = *d*(*u*, 1 − *s*) − *d*^*il*^(*u*, 1 − *s*).

For the lower bound, let *Z*′ be the set of all vertex-sides (*u*′, *s*′) such that (*u*′, *s*′)≠ (*u*, 1 − *s*) and there is an edge {(*u*, 1 − *s*), (*u*′, *s*′)}. Let *Z* = {*z* | *F*_*V*_ (*z*) ∈ *Z*′}. By Lemma B.10, |*Z*| = |*Z*′|. By Lemma B.11, for every *z* ∈ *Z*, there is an edge from *x* to *z* in *G*_dbl_(*K*) and therefore *d*^+^(*x*) ≥ |*Z*| = |*Z*′|.

Now we show that *d*(*u*, 1 − *s*) ≤ *d*^+^(*x*) + *d*^*il*^(*u*, 1 − *s*) by considering two cases. In the first case, assume that there is no inverted loop touching (*u*, 1 −*s*). Then, *d*(*u*, 1 −*s*) = |*Z*′| and *d*^*il*^(*u*, 1 −*s*) = 0. We can therefore write *d*(*u*, 1 − *s*) = |*Z*′| + *d*^*il*^(*u*, 1 − *s*) ≤ *d*^+^(*x*) + *d*^*il*^(*u*, 1 − *s*). In the second case, assume there exists an inverted loop touching (*u*, 1 − *s*). In this case, *d*(*u*, 1 − *s*) = |*Z*′| + 2. By Lemma B.11, there is an edge from *x* to 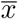 and 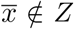. Thus, *d* + (*x*) ≥ |*Z*| + 1. Putting this together, *d*(*u*, 1 − *s*) = |*Z*′| + 2 = |*Z*| + 2 ≤ *d*^+^(*x*) + 1 = *d*^+^(*x*) + *d*^*il*^(*u*, 1 − *s*).

For part (*ii*), observe that 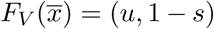. We can then apply part (*i*) of this theorem to 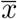, *u*, and 1 − *s*, and get that 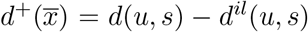. By Lemma B.3, 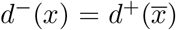, and hence 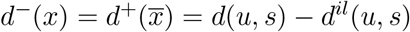. □

An immediate consequence of the degree-preserving lemma is that if *F* (*w*) is a unitig, then so is *w*. The converse is not always true however.

#### Lemma B.14.

*Let K be a set of canonical k-mers where k is odd. Let w* = (*x*_0_, …, *x*_*n*_) *and t* = (*u*_0_, *s*_0_, …, *u*_*n*_, *s*_*n*_) *be two walks related by* 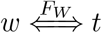.

i. *If t is a unitig, then w is a unitig*.
ii. *If w is a unitig and for all* 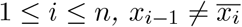, *then t is a unitig*.

*Proof*. For (*i*), when *n* = 0, *w* is trivially a unitig because it has only one vertex. For *n >* 0, since *t* is a unitig, *d*(*u*_*i*_, *s*_*i*_) = 1 for 0 *< i* ≤ *n*. Moreover, since an inverted loop would make a degree ≥ 2, we have *d*^*il*^(*u*_*i*_, *s*_*i*_) = 0. Using Lemma B.13, *d*^−^(*x*_*i*_) = 1. Similarly, for all 0 ≤ *i < n, d*(*u*_*i*_, 1 − *s*_*i*_) = 1, *d*^*il*^(*u*_*i*_, 1 − *s*_*i*_) = 0, and Lemma B.13 gives that *d*^+^(*x*_*i*_) = 1. Hence *w* is a unitig.

For (*ii*), first observe that there is no inverted loop incident to (*u*_*i*_, *s*_*i*_), for 1 ≤ *i* ≤ *n*. If that were the case, then Lemma B.11 implies that there is an edge from 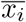 to *x*_*i*_. Since *w* is a unitig, the only in-neighbor of *x*_*i*_ is *x*_*i* −1_. Hence, 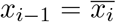, which contradicts the conditions of the Lemma. Now, since *d*^*il*^(*u*_*i*_, *s*_*i*_) = 0, Lemma B.13 implies that *d*(*u*_*i*_, *s*_*i*_) = *d*^−^(*x*_*i*_) + *d*^*il*^(*u*_*i*_, *s*_*i*_) = *d*^−^(*x*_*i*_) = 1. Using a symmetrical argument (omitted), *d*(*u*_*j*_, 1 − *s*_*j*_) = 1 for all 0 ≤ *j < n*. Therefore, *t* is a unitig. □

Similarly, we can relate the maximality of unitigs in *G*_dbl_(*K*) and *G*_bid_(*K*). A maximal unitig in *G*_dbl_(*K*) is maximal in *G*_bid_(*K*), on the condition that is a unitig in *G*_bid_(*K*); however, the other direction only holds with a restrictive condition.

#### Lemma B.15.

*Let K be a set of canonical k-mers where k is odd. Let w* = (*x*_0_, …, *x*_*n*_) *and t* = (*u*_0_, *s*_0_, …, *u*_*n*_, *s*_*n*_) *be two walks related by w* 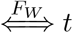. *Suppose that both w and t are unitigs*.

i. *If t is prefix-maximal and has no lonely inverted loop at the first endpoint side, then w is prefix-maximal*.
ii. *If w is prefix-maximal, then t is prefix-maximal*.
iii. *If t is suffix-maximal and has no lonely inverted loop at the last endpoint side, then w is suffix-maximal*.
iv. *If w is suffix-maximal, then t is suffix-maximal*.

*Proof*. We will prove (*i*) and (*ii*) only, since the proofs of (*iii*) and (*iv*) are symmetric. For (*i*), if there is more than one edge incident to (*u*_0_, *s*_0_), then *d*(*u*_0_, *s*_0_) ≥ 2. If there are no edges incident to (*u*_0_, *s*_0_), then *d*(*u*_0_, *s*_0_) = 0. In both cases, Lemma B.13 implies that *d*^−^(*x*_0_) = *d*(*u*_0_, *s*_0_)≠ 1 and Lemma B.4 implies that *w* is prefix-maximal.

Now consider the case that *d*(*u*_0_, *s*_0_) = 1. By the conditions of the Lemma, there is no inverted loop incident at (*u*_0_, *s*_0_), and Lemma B.13 implies *d*^−^(*x*_0_) = 1. Since *t* is prefix-maximal, by Lemma B.7, there is a vertex side (*u*′, *s*′) and an edge *e* = (*u*′, *s*′), (*u*_0_, *s*_0_) such that *d*(*u*′, *s*′) *>* 1. Let 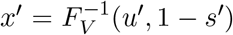 and Lemma B.11 implies that there is an edge from *x*′ to *x*_0_ in *G*_dbl_(*K*). Observe that because *d*(*u*_0_, *s*_0_) *<* 2, *e* is not an inverted loop. Therefore, (*u*′, *s*′) has at least one incident edge that is not an inverted loop. Because an inverted loop adds at least two to the degree, *d*(*u*′, *s*′) − *d*^*il*^(*u*′, *s*′) *>* 1. Thus, Lemma B.13 implies that *d*^+^(*x*′) *>* 1. By Lemma B.4, *w* is a prefix-maximal unitig.

For (*ii*), suppose for the sake of contradiction that *t* is not prefix-maximal. Then Lemma B.7 implies that *d*(*u*_0_, *s*_0_) = 1 and there exists a vertex-side (*u*′, *s*′) with *d*(*u*′, *s*′) = 1 and an edge *e* = {(*u*′, *s*′), (*u*_0_, *s*_0_)}. Let 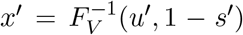 Note that *d*^*il*^(*u*_0_, *s*_0_) = *d*^*il*^(*u*′, *s*′) = 0 because vertex-sides with degree 1 cannot have an inverted loop incident to them. Lemma B.13 then implies that *d*^−^(*x*_0_) = *d*(*u*_0_, *s*_0_) = 1 and *d*^+^(*x*′) = *d*(*u*′, *s*′) = 1. In addition, Lemma B.11 applied to *e* says that there is an edge from *x*′ to *x*. By Lemma B.4, these facts imply that *w* is not prefix-maximal, which is a contradiction. □

Theorem 2 has a condition that there are no circular unitigs. We now show that this implies that a unitig in *G*_bid_(*K*) cannot have lonely inverted loops incident to both of the endpoint sides.

#### Lemma B.16.

*Let K be a set of canonical k-mers where k is odd. Let w* = (*x*_0_, …, *x*_*n*_) *be a walk in G*_*dbl*_(*K*) *such that F*_*W*_ (*w*) *is a unitig. If the two endpoint sides of F*_*W*_ (*w*) *have lonely inverted loops incident on them, then* 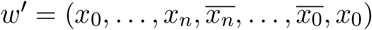 *is a circular unitig in G*_*dbl*_(*K*).

*Proof*. First, to show that *w*′ is a walk in *G*_dbl_(*K*), we need to show that there exist edges 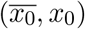 and 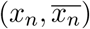. This follows by applying Lemma B.11 to the inverted loop edges at the endpoints of *F* (*w*), i.e. to {(*u*_0_, *s*_0_), (*u*_0_, *s*_0_)} and {(*u*_*n*_, 1 − *s*_*n*_), (*u*_*n*_, 1 − *s*_*n*_)}.

Second, to show that *w* is a unitig, we will show that all the necessary vertex degrees are 1. By Lemma B.14, *w* is a unitig, and hence *d*^+^(*x*_*i*_) = 1 for all 0 ≤ *i < n* and *d*^−^(*x*_*i*_) = 1 for all 0 *< i* ≤ *n*. Let (*u*_*i*_, *s*_*i*_) = *F*_*V*_ (*x*_*i*_) for all 0 ≤ *i* ≤ *n*. Because the endpoint sides of *F* (*w*) each have a lonely inverted loop, *d*(*u*_0_, *s*_0_) = 2 and *d*(*u*_*n*_, 1 − *s*_*n*_) = 2. Applying Lemma B.13, *d*^−^(*x*_0_) = *d*(*u*_0_, *s*_0_) − *d*^*il*^(*u*_0_, *s*_0_) = 2 − 1 = 1 and *d*^+^(*x*_*n*_) = *d*(*u*_*n*_, 1 − *s*_*n*_) − *d*^*il*^(*u*_*n*_, 1 − *s*_*n*_) = 1. Applying Lemma B.3 to all these, we get that 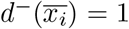 for all 0 ≤ *i* ≤ *n* and 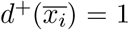 for all 0 ≤ *i* ≤ *n*. □

### B.4 Proof of Theorem 2

#### Theorem 2.

*Proof*.

i. We already know from Lemma B.12 that *F*_*W*_ is a bijection between walks in *G*_dbl_(*K*) and *G*_bid_(*K*). It remains to show that

1. For a unitig *w* that is maximal and non-palindromic in *G*_dbl_(*K*), *F*_*W*_ (*w*) ∈ *B*_no-loop_.
2. For a unitig *t* ∈ *B*_no-loop_, *F* ^−1^(*t*) is a maximal and non-palindromic unitig in *G*_dbl_(*K*). First, we prove (1). Because *w* is a non-palindromic maximal unitig, by Lemma B.6, there is no edge 0 ≤ *i < n* such that 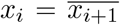, because then (*x*_*i*_, *x*_*i*+1_) would be a palindromic sub-unitig of *w*. Hence we can apply Lemma B.14 to say that *F*_*W*_ (*w*) is a unitig and we can apply Lemma B.15 to say that *F*_*W*_ (*w*) is maximal. Hence *F*_*W*_ (*w*) ∈ *B*. To show that *F*_*W*_ (*w*) ∉ *B*_2_ ∩ *B*_3_, first assume for the sake of contradiction that there is a lonely inverted loop at the last endpoint side of *F*_*W*_ (*w*). Then by Lemma B.11 there is an edge from *x*_*n*_ to 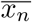. By Lemma B.13, *d*^+^(*x*_*n*_) = 2 − 1 = 1. By Lemma B.3, 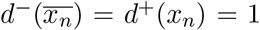. Because *w* is maximal, if *d*^+^(*x*_*n*_) = 1, then 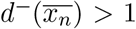. This is a contradiction. The argument that there is no lonely inverted loop at the first endpoint side of *F*_*W*_ (*w*) is symmetric and omitted. Now, we prove (2). Let *w* = *F* ^−1^(*t*). Since *t* is a unitig, Lemma B.14 implies that *w* is a unitig also. Moreover, Lemma B.9 implies that *t* is non-palindromic; since *F*_*W*_ is spelling preserving (Lemma B.12), *w* is also non-palindromic. Since the Theorem assumes that *G*_dbl_(*K*) does not have circular unitigs, Lemma B.16 implies that *t* cannot have a lonely inverted loop at both endpoints. Since *t* ∉ *B*_2_ ∪ *B*_3_, it also cannot have an inverted loop at exactly one endpoint. We can therefore apply Lemma B.15 to get that *w* is maximal.
ii. Observe that *rev* is by definition a function that is its own inverse and is a bijection on the set of walks in *G*_bid_(*K*). Furthermore, Lemma B.8 implies that *rev* remains a bijection when restricted to maximal unitigs in *G*_bid_(*K*). Finally, observe that for a walk *t*, the first (respectively, last) endpoint side of *t* is the last (respectively, first) endpoint side of *rev*(*t*). These facts together imply that *rev* is a bijection between *B*_first-loop_ and *B*_last-loop_.
iii. To show that head is a bijection we show
  1. for all *w* ∈ *D*_pal_, head(*w*) ∈ *B*_last-loop_,
  2. for all *t* ∈ *B*_last-loop_, there exists a *w* ∈ *D*_pal_ such that head(*w*) ∈ *B*_last-loop_.
  3. the above *w* is unique.
  First, we prove (1). Let *w* = (*x*_0_, …, *x*_*n*_). By Lemma B.1, *n* is odd and at least 1. Let *m* = (*n* − 1)*/*2 and let *h* ≜ (*x*_0_, …, *x*_*m*_). Since *w* is a palindromic unitig and, by the conditions of the Theorem, non-circular, Lemma B.5 implies that for all 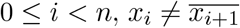. Then by Lemma B.14, head(*w*) = *F*_*W*_ (*h*) is a unitig. Simultaneously, because *w* is a maximal unitig, *h* is a prefix-maximal unitig. Lemma B.15 then implies that *F*_*W*_ (*h*) is prefix-maximal. Now we show that *F*_*W*_ (*h*) is suffix-maximal and has a lonely inverted loop at the last endpoint. Let (*u*_0_, *s*_0_, …, *u*_*m*_, *s*_*m*_) ≜ *F*_*W*_ (*h*). Since *w* is palindromic, Lemma B.5 implies that 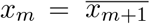, and, hence, *u*_*m*_ = *u*_*m*+1_. By Lemma B.11, there is an inverted loop incident to (*u*_*m*_, 1 − *s*_*m*_), i.e. the last endpoint of *F*_*W*_ (*h*). Because *w* is a unitig, *d*^+^(*x*_*m*_) = *d*^−^(*x*_*m*+1_) = 1, Lemma B.13 then implies that *d*(*u*_*m*_, 1 − *s*_*m*_) = *d*^+^(*x*_*m*_) + *d*^*il*^(*u*_*m*_, 1 − *s*_*m*_) = 2. By Lemma B.7, *F*_*W*_ (*h*) is suffix-maximal and therefore we have shown that *F*_*W*_ (*h*) ∈ *B*_last-loop_. Next we prove (2). Let (*u*_0_, *s*_0_, …, *u*_*n*_, *s*_*n*_) = *t* and let 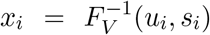. Let 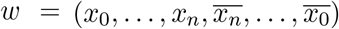 be a sequence of vertices in *G*_dbl_(*K*). We will first show that *w* is a walk, then that it is palindromic, then that it is a unitig, and finally that it is maximal. Note that *w* is equivalently defined to be the concatenation of 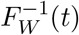 with 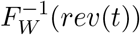. Applying Lemma B.12, the sequences (*x*_0_, …, *x*_*n*_) and 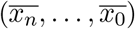 are walks. Since *t* is in *B*_last-loop_, there is an inverted loop incident to (*u*_*n*_, 1 − *s*_*n*_). By Lemma B.11, this implies there is an edge from *x*_*n*_ to 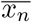 in *G*_dbl_(*K*). Therefore, *w* is a walk. It is palindromic by its definition. Since *t* is a unitig, by Lemma B.8, *rev*(*t*) is a unitig. Now applying Lemma B.14, *w* and *rev*(*w*) are both unitigs. Because the inverted loop is lonely, *d*(*u*_*n*_, 1 − *s*_*n*_) = 2, and by Lemma B.13, *d*^+^(*x*_*n*_) = 1. Applying Lemma B.3, 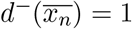. Hence *w* is a unitig. As *t* is in *B*_last-loop_, this implies that no lonely inverted loop is incident to (*u*_0_, *s*_0_). We can apply Lemma B.15 to get that *F* ^−1^(*t*) is prefix-maximal. Because *w* starts with *F* ^−1^(*t*), *w* is also prefix-maximal. By Lemma B.8, *F* ^−1^(*rev*(*w*)) is suffix-maximal. Because *w* ends with *F* ^−1^(*rev*(*w*)), *w* is also suffix-maximal. Hence, *w* is maximal. For (3), let (*u*_0_, *s*_0_, …, *u*_*n*_, *s*_*n*_) = *t* and let 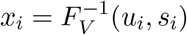). Let *w*′ be a walk in *D*_pal_ such that head(*w*′) ∈ *B*_last-loop_. We will show that 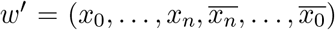. Since head(*w*′) has *n* + 1 vertices, *w* must have 2*n* + 2 vertices. Hence we can write 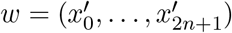. Since *w* is a palindrome, we have that 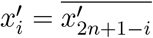 for all 0 ≤ *i* ≤ 2*n* + 1. We can therefore rewrite *w* as 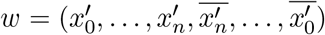. Next, observe that 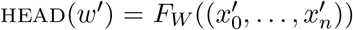. Since this must be equal to *t* and *F*_*W*_ is a bijection (Lemma B.12), we get that 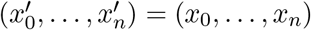. We can therefore rewrite *w* as 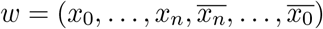, which is the same as *w*. □

## C Experimental details

### Choice of *k* parameter for the assemblers

To ensure that the results across the assemblers are comparable, we set the *k* parameter in a way so that the set of unitigs constructed are as close as possible. The ideal way is to set *k* such that the underlying *k*-mer sets *K* used for all assemblers are same. However, there was a practical limitation for that. We note that both SPAdes and MEGAHIT are a multi-*k* assemblers, so the *k* parameter is just the maximum allowed *k*-mer size. When we pass the value *k* to the assemblers, both SPAdes and MEGAHIT use *k*-mer set and (*k* + 1)-mer set to construct unitigs, whereas bcalm, ABySS, and minia uses a node-centric de Bruijn graph with only *k*-mer sets as vertices. As such, we found that the output unitigs of SPAdes and MEGAHIT with a value of *k* are more similar to unitigs of bcalm and ABySS created with *k* + 1. We also note that SPAdes and MEGAHIT only allow odd *k*, which is why we needed to use an even *k* for *G*_dbl_.

In Table 3, we therefore passed *k* = 74 to bcalm and *k* = 73 to SPAdes and MEGAHIT. Since Theorem 1 is valid for all *k*, this was not an issue for Table 3. We used the default parameter for minimum *k*-mer coverage for both assemblers.

For Table 6, we passed *k* = 31 to all assemblers, since Theorem 2 only applies when the vertex lengths are of odd *k*. Since SPAdes and MEGAHIT by default use both *k*-mer and (*k* + 1)-mer set to construct unitigs, the number of palindromic unitigs (433) differs from the number in minia and ABySS (440). However, this is not a problem because we are not comparing the numbers between assemblers but only within assemblers.

### Detection of palindrome splitting artifact

In this section, we use the notation *S*[*i* : *j*] to denote substring of string *S* starting at index *i* and ending at index *j*. Let *w* = (*x*_0_, …, *x*_*n*_) be a palindromic unitig in *D*_pal_ and let *p* be its spelling. We say a unitig in *D*_pal_ is *fully-covered* if there exists some contig that aligns to an interval which contains *p*’s interval in the reference. Let *k*′ ≜ (*k* − 1)*/*2. We say *w* is *split* if there exists at least one contig *c* such that either

1. *c* aligns to an interval that starts before *p*’s interval and ends exactly at position |*p*|*/*2 + *k*′ of *p*’s interval and there are no other contigs with alignments intersecting *p*[|*p*|*/*2 + *k*′ + 1 : |*p*|], or
2. *c* aligns to an interval that ends after *p*’s interval and starts exactly at location |*p*|*/*2 − *k*′ + 1 of *p*’s interval and there are no other contigs with alignments intersection *p*[1 : |*p*|*/*2 + *k*′].

We say *w* is *ambiguous* if it does not fall into either category.

To motivate these cases, observe that the length of *p* is *n* + *k* and, because *p* is a palindrome and *k* is odd, *n* must be odd. Let 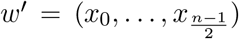 be the first half of the walk *w* and let *p*′ be its spelling. By Theorem 2, head(*w*) ∈ *B*_last-loop_ and *rev*(head(*w*)) ∈ *B*_first-loop_. Then, 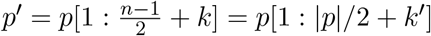. Then,

1. *spell*(head(*w*)) = *spell*(*F*_*W*_ (*w*′)) = *spell*(*w*′) = *p*[1 : |*p*|*/*2 + *k*′], and
2. 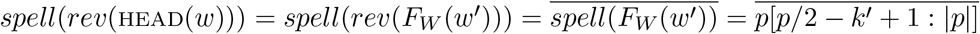.

The cases we describe therefore correspond to observing the alignments of head(*w*) and *rev*(head(*w*)) to the corresponding places of *p* and not observing any other bidirected unitigs aligning across the middle boundaries.

### CAMI dataset

We used the benchmark called “low complexity dataset” in [31]. Since our analysis requires error-free reads, we re-simulated the reads using identical genomes and abundances (as detailed in supplementary materials of [26]). Table S1 shows the properties and relative abundances of the genomes. We used CAMISIM [13] for the simulations, with read length of 150nt and insert size 150.

**Table S1:**
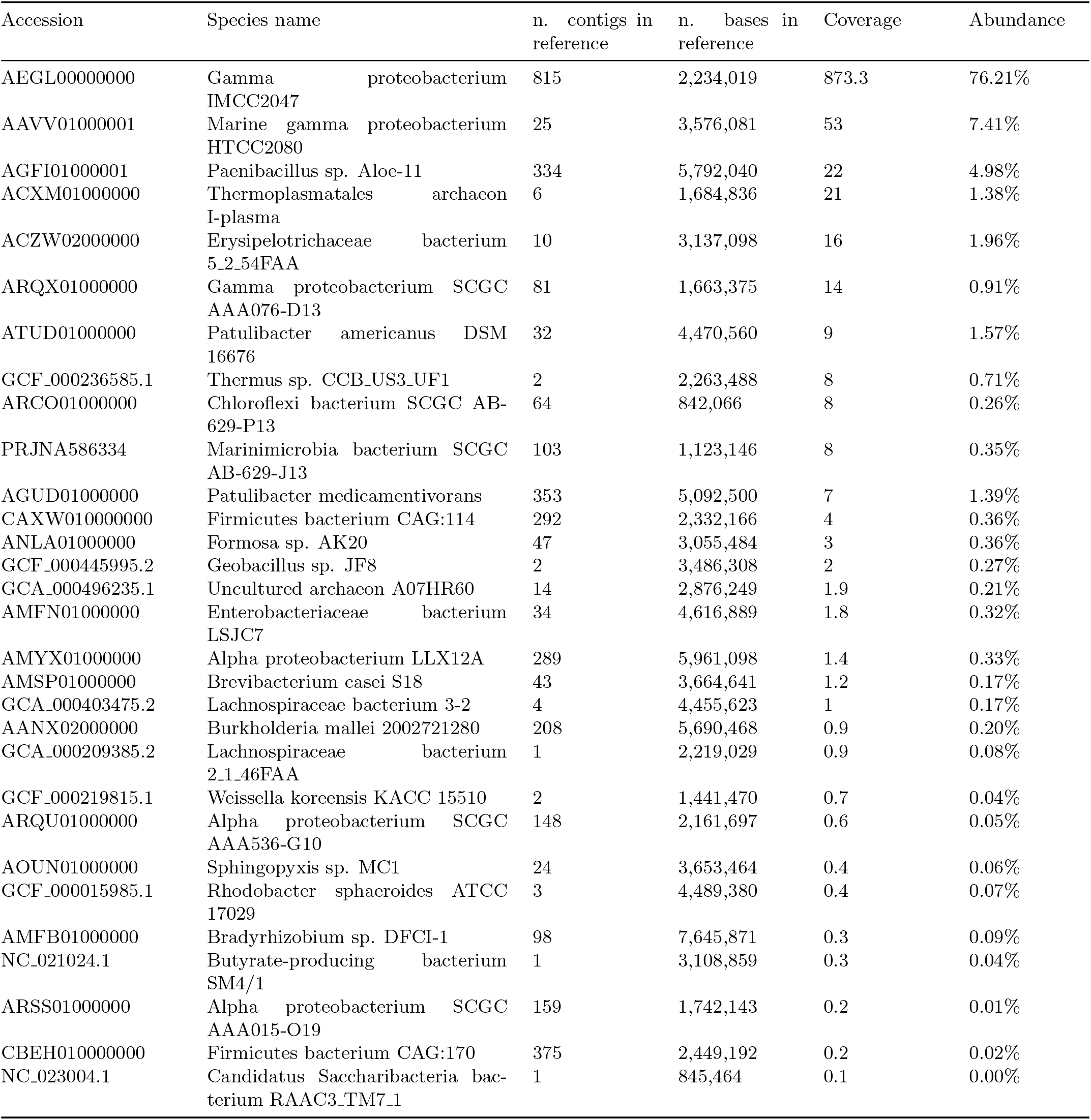
Characteristics of all 30 genomes consituting the CAMI “low complexity” dataset. The coverage refers to the depth-of-coverage for each genome, in both the benchmark ([26]) and our simulations.

## Footnotes

1 The safety of unitigs has been previously studied for other notions of “safety” by [9]. While the authors did not make the explicit conclusion and did not verify it in practice, their Theorem 6.1(d) implies that unitigs are not guaranteed to be safe in the model of assembly they consider. Concretely, while a suffix or prefix of the unitig may be present at the starts and ends of parts of the genome, the whole unitig might never be contained as a contiguous sequence.

2 As an aside, we mention how one would obtain a bijection. This is not necessary for the proofs of this paper, but may be a useful observation in its own right. Let *E* be the set of edges in *G*_dbl_(*K*), let *α* ⊆ *E* be all the self-mirror edges, and let *β* be the partition of *E \ α* into mirror edge-pairs. For example, if *E* = {(*AGG, GGA*), (*T CC, CCT*), (*T TA, TAA*)}, then *α* = {(*T TA, TAA*)} and *β* = {{(*AGG, GGA*), (*T CC, CCT*)}}. For an element 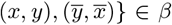, we define 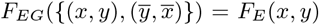. For a self-mirror edge (*x, y*) ∈ *α*, we define *F*_*EG*_({(*x, y*)}) = *F*_*E*_(*x, y*). One can then show that *F*_*EG*_ is a bijection between *α* ∪ *β* and edges in *G*_bid_(*K*).

